# Oncogenic E3-ligase adaptors MAGE-A3/6 promote cancer cell migration via BAP18 degradation

**DOI:** 10.64898/2026.03.23.713706

**Authors:** Maximilian W.G. Schneider, Marie S. Polgar, Robert W. Kalis, Philip Barbulescu, Natalia Brunner, Mathias Madalinski, Dalia Barsyte-Lovejoy, Johannes Zuber, Manfred Koegl, Ralph A. Neumüller, Paola Martinelli

## Abstract

Cancer testis antigens are widely expressed in human malignancies. Melanoma-Associated Antigens (MAGE) A3 and A6 have been proposed to modulate protein turnover and metabolism in cancer cells. However, the substrate specificity of MAGE-A3/6 and the impact on cancer cell behavior remain poorly understood. Although previous research has identified binding partners, a molecularly validated target for MAGE-A3/6-mediated proteasomal degradation has not been described. In this study, we redefine the substrate specificity of MAGE-A3/6 and present a mechanistic framework for substrate binding, polyubiquitination, and subsequent degradation. We identify BPTF-Associated Protein of 18kDa (BAP18) as a bona fide novel substrate of MAGE-A3/6 and demonstrate its direct regulation via a molecularly defined substrate-degron-E3-adaptor interaction. The degradation of BAP18 by MAGE-A3/6 underlies phenotypic alterations in cancer cells, such as enhanced migratory capacity. This previously unrecognized molecular link is observed in both cancer cell lines and human cancer tissues, supporting a role as a fundamental oncogenic process. The discovery of a molecularly defined interaction between MAGE-A3/6 and their substrate enables systematic investigation into oncogenic protein degradation in human cancers and may inform future therapeutic strategies that leverage the molecular function of aberrantly reexpressed germline proteins in cancer.

## Main text

Under physiological conditions, expression of cancer testis antigens (CTAs) is restricted to reproductive tissues. In a variety of human malignancies, however, CTAs are aberrantly reexpressed^1–3^. Although CTAs have been extensively studied as clinical biomarkers and potential targets for immunotherapy, their contribution to cancer initiation and progression is poorly understood. Amongst CTAs, the MAGE-A family has been investigated in the context of several cancer types. The expression of MAGE-A proteins is associated with poor prognosis, metastasis, immune evasion and cancer aggressiveness^4,5^ and might confer reactivation of developmental processes during cancer progression^6^. Due to their cancer specific expression, MAGE-As have been used as therapeutic targets and MAGE-A-targeting agents have reached the clinic, including MAGE-A3 targeted mRNA vaccines^7,8^ and a recently approved autologous T-cell immunotherapy against MAGE-A4^9^. Additionally, MAGEs were proposed to act as E3 ligase substrate adaptors, promoting degradation of target proteins^10^. However, molecularly validated substrates enabling mechanistic and phenotypic studies remain elusive. The highly similar paralogs MAGE-A3 and MAGE-A6^4,10,11^ were previously proposed to bind TRIM28 and mediate polyubiquitination of substrates including AMPKα1^4^ and FBP1^12^, both linked to metabolic reprogramming, as well as ALKBH2^12^ and p53^10,13,14^, proteins linked to stress response and DNA damage. However, a molecularly defined framework for substrate binding and degradation has not been established. Additionally, the phenotypic consequences of MAGE-A3/6 expression are poorly understood, and thus a mechanistic link of degraded substrates to clinical phenotypes remains elusive.

### MAGE-A3 targets BAP18 for degradation

To identify novel MAGE-A3 substrates, we generated a doxycycline-inducible MAGE-A3-expressing cell line in a MAGE-A-negative background (DLD-1-iMAGE-A3). The colorectal adenocarcinoma (COAD) cell line DLD-1 has no detectable expression of any MAGE-A proteins (Extended Data Fig 1a) but expresses robust levels of TRIM28, the cognate E3 ligase for MAGE-A3/6 (Extended Data Fig 1b). Dose-dependent MAGE-A3 induction was validated in DLD-1-iMAGE-A3 cells using immunofluorescence (Extended Data Fig 1c).

DLD-1-iMAGE-A3 cells were treated with doxycycline over 40 hours and changes in the proteome were assessed using whole proteome quantitative mass spectrometry (qMS). MAGE-A3 expression was robustly induced, while the protein most significantly reduced was BPTF-associated protein of 18 kDa (BAP18), which was downregulated in a doxycycline- and thus MAGE-A3-dose-dependent manner (Fig 1a-c, Extended Data Fig 1d). To determine whether MAGE-A3-dependent BAP18 regulation is a direct effect, we quantified BAP18 levels in a time course upon doxycycline treatment for up to 48h (Fig 1d, Extended Data Fig 1e). MAGE-A3 expression induced a rapid and continued reduction of BAP18 protein levels already after 4 hours and until the last measurement, consistent with a direct modulation. Accordingly, BAP18 degradation was readily reversible upon doxycycline washout (Fig 1e, Extended Data Fig 1f). Sixteen hours after removal of doxycycline, MAGE-A3 protein levels decreased by ∼90%, consistent with a published half-life of ∼4.5 hours^15^. BAP18 protein levels increased concurrently, with a resynthesis halftime of ∼21 hours, similar to the reported estimated resynthesis rate determined by SILAC^16^. The immediate recovery of BAP18 levels after MAGE-A3 removal supports direct and post-translational modulation of BAP18 through MAGE-A3.

**Figure 1.**
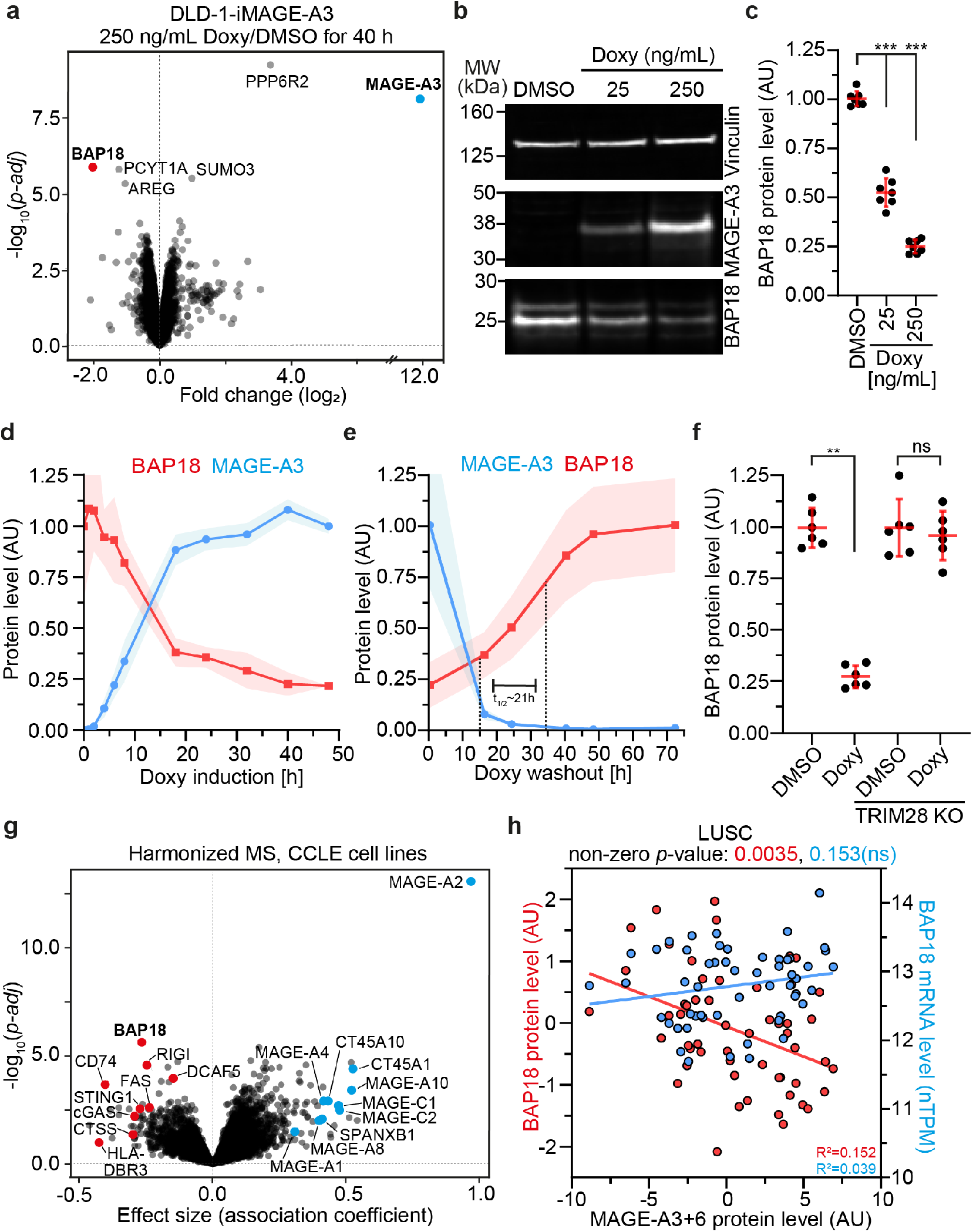
BAP18 is a novel substrate for MAGE-A3/6 induced degradation by the UPS. **a**, Volcano plot displaying qMS analysis of DLD-1-iMAGE-A3 cells after 40h treatment with 250 ng/mL doxycycline compared to DMSO. Top up- and downregulated proteins indicated in bold. **b**, Fluorescent immunoblot analysis of MAGE-A3 and BAP18 protein levels in DLD-1-iMAGE-A3 cells treated with DMSO, 25 ng/mL or 250 ng/mL doxycycline. **c**, Quantification of BAP18 protein level in the analysis shown in panel b. Data are mean ± s.d. Significance was tested by a two-tailed Mann-Whitney *U*-test (25 ng/mL doxycycline, *P* = 5.83^-4^; 250 ng/mL doxycycline, *P* = 5.83^-4^). **d, e**, Simple western quantification of BAP18 and MAGE-A3 protein levels during time course after doxycycline induction (d) and washout (e). Data are mean ± s.d. **f**, Simple western quantification of BAP18 protein levels after TRIM28 KO in DLD-1-iMAGE-A3 cells after treatment with DMSO or 250 ng/mL doxycycline. Significance was tested with a two-tailed Mann-Whitney *U*-test (parental, *P*= 2.16^-3^; TRIM28 KO, *P*=0.94). **g**, Regression analysis of every protein against MAGE-A3/6 expression in proteomics data from 375 CCLE cell lines. **h**, Regression analysis of harmonized BAP18 protein and mRNA expression against MAGE-A3/6 harmonized protein expression in LUSC tumor samples. *P*-values indicate significant/non-significant deviation of the linear regression line slope from zero. *n* = 54 tumors for which matched MAGE-A3/6 and BAP18 protein and mRNA expression are available. Biological replicates: *n* = 3 (**a, d, e**), *n* = 7 (**b, c**), *n* = 6 (**f**). Technical replicates: *n* = 1-2.

As TRIM28 is the proposed cognate E3 ligase for MAGE-A3^4,10^, we tested its requirement for MAGE-A3-dependent BAP18 degradation. TRIM28 knockout (KO), viable in DLD-1 cells, fully abrogated MAGE-A3-dependent downregulation of BAP18 protein in DLD-1-iMAGE-A3 cells (Fig 1f, Extended Data Fig 2a, b). Furthermore, treatment of DLD-1-iMAGE-A3 cells with the proteasome inhibitor MG132 for 12h in addition to doxycycline prevented the BAP18 degradation induced by MAGE-A3 expression (Extended Data Fig 2c, d). Altogether, these data confirm that BAP18 degradation by MAGE-A3 is mediated by TRIM28 and the ubiquitin-proteasome system (UPS).

Since MAGE-A3 and MAGE-A6 share 96% sequence similarity, we tested for functional redundancy in UPS-dependent BAP18 degradation. We generated MAGE-A6 inducible DLD-1 cells (DLD-1-iMAGE-A6) and verified doxycycline mediated MAGE-A6 expression by immunofluorescence (Extended Data Fig 2e). Due to their high similarity, the same antibody was used to detect inducible MAGE-A3 and MAGE-A6, allowing for direct comparison of the expression level in DLD-1-iMAGE-A3 and DLD-1-iMAGE-A6, which was almost identical (Extended Data Fig 2f). Treatment of DLD-1-iMAGE-A6 with 250 ng/mL doxycycline for different times up to 48 hours resulted in reduction of BAP18 protein reminiscent to DLD-1-iMAGE-A3 cells (Extended Data Fig 2g, h), indicating functional redundancy between the MAGE-A3/6 paralogs. To test if MAGE-A3/6 dependent regulation of BAP18 can be generalized, we analyzed BAP18 and MAGE-A3/6 protein levels in a panel of 15 cancer cell lines comprising different cancer types. Consistent with our findings in DLD-1 cells, BAP18 and MAGE-A3/6 levels showed statistically significant anticorrelation across the panel (Extended Data Fig 3a, b). This anticorrelation was not observed between MAGEA3/6 protein and BAP18 mRNA level, supporting post-translational regulation (Extended Data Fig 3c). Furthermore, BAP18 was the top MAGE-A3/6 negatively associated protein across a proteomic dataset containing 375 CCLE cell lines^17^, suggesting that BAP18 downregulation by MAGE-A3/6 is a general regulatory mechanism (Fig 1g). The expression of other MAGE-A family members (A1, A2, A4, A8, A10) as well as other CTAs (CT45A1, CT45A10, SPANXB1) was positively associated with MAGE-A3/6, supporting the notion that CTAs are often coexpressed due to their overlapping reactivation mechanism^1,2^, which is thought to occur through DNA hypomethylation^1,18^. Additional proteins strongly negatively associated with MAGE-A3/6 expression included components of the antigen processing- and presentation machinery (CD74, DCAF5, Cathepsin S, HLA-DBR3) as well as innate immune and cellfate modulating signaling proteins (STING1, cGAS, RIG-I, FAS). This pattern aligns with the broader observation that CTA-high tumors often evade immune detection by simultaneously reducing antigen presentation and dampening responses to interferon-stimulating signals^19,20^. There is currently no evidence that these proteins are direct substrates of MAGE-A3/6.

To test for anticorrelation of MAGE-A3/6 and BAP18 in patient samples, we then queried published proteogenomic datasets for lung squamous cell carcinoma (LUSC)^21^ and treatment naïve metastatic malignant melanoma (MMM)^22^. Analysis of aggregated MAGE-A3 and -A6 versus BAP18 protein expression identified an anticorrelation akin to cell lines (Fig 1h, Extended Data Fig 3d). Consistent with post-translational regulation, the anticorrelation was not found for BAP18 mRNA and MAGE-A3/6 protein expression. Altogether, these data establish BAP18 as a novel MAGE-A3/6 substrate across human malignancies. Of note, previously proposed MAGE-A3/6 substrates were not identified in our analyses (Extended Data Fig 4a-j).

### MAGE-A3 directly binds to BAP18

To map the interaction between both proteins, we designed a peptide array covering BAP18 in a series of overlapping 15-amino-acid segments staggered by one residue. The array was incubated with recombinant full-length MAGE-A3, and sites of interaction were detected using an anti–MAGE-A3 antibody (Extended Data Fig 5a). A high-affinity interaction region near the N-terminus of BAP18, located within a predicted helical segment, was identified (Fig 2a). Sequence alignment of the interacting peptides revealed an enrichment of hydrophobic residues, arranged with alternating polar and non-polar amino acids, forming an amphipathic helical peptide (Extended Data Fig 5b, c).

**Figure 2.**
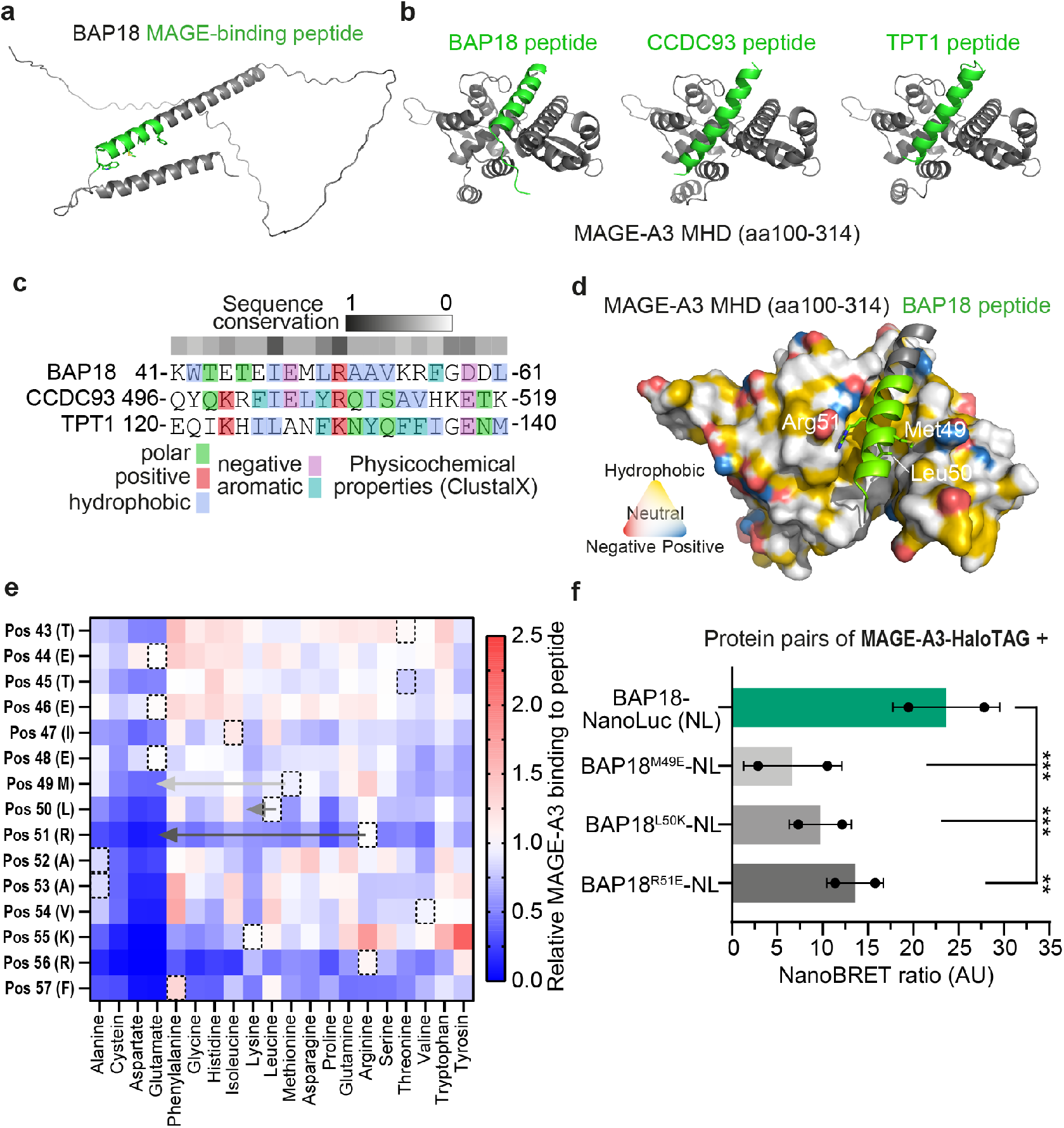
BAP18 binds directly to MAGE-A3 through an amphiphilic helical peptide. **a**, AF3 prediction of BAP18 structure (AF-Q8IXM2-F1, canonical isoform of 172 amino acids), highlighting the helix-loop-helix core structural element. Identified MAGE-A3 interacting peptide within central helix shown in light green. **b**, AF3 *in silico* modeling of MAGE-A3 MHD together with various peptides (light green) determined to engage with MAGE-A3. **c**, MUSCLE peptide sequence alignment of peptides derived from BAP18, CCDC93 and TPT1, with amino acid position in the full-length protein. Indicated physicochemical properties are according to ClustalX scale. **d**, Surface representation of AF3 multimer MAGE-A3 MHD, with YRB false color indicating charge and hydrophobicity, with BAP18 derived peptide occupying the hydrophobic SBC. **e**, Heatmap depicting the detected fluorescence signal from MAGE-A3 for a given peptide with mutations to the indicated amino acid. Amino acids present in the wild type sequence are outlined by dashed boxes. Data is normalized to the mean of all reference wild-type peptide fluorescence intensities. Arrows indicate single mutations included in the following NanoBRET assay. **f**, Quantification of NanoBRET cellular interaction assay of MAGE-A3-HaloTAG with full-length BAP18-Nanoluciferase bearing the indicated mutations. Bars indicate mean ± s.d. of *n* = 2 biological replicates (2 technical replicates each). Significance was tested using Dunnett’s multiple comparisons test (*P* = 1.51^-4^ (BAP18^M49E^); *P* = 8.51^-4^ (BAP18^L50K^); *P* = 9.41^-3^ (BAP18^R51E^)).

To expand on the identified MAGE-A3-interacting peptide, we performed a yeast two-hybrid (Y2H) screen^23^. The MAGE-A3 Mage Homology Domain (MHD; amino acids 100–314) was used as bait, and a human lung cancer cDNA library served as prey. The screen identified multiple protein fragments that interacted with MAGE-A3 (Extended Data Fig 5d). Each Y2H-derived hit contained regions of strong binding in a peptide array (Extended Data Fig 5e), and alignment of these peptides revealed an enrichment of hydrophobic residues, similar to the BAP18 peptide (Extended Data Fig 5f).

In vitro pulldowns with peptides derived from BAP18, CCDC93, and TPT1 confirmed robust interaction with full-length MAGE-A3 compared to a random sequence control (Extended Data Fig 5g). The MAGE-A3 MHD (aa 100–314) yielded similar results, verifying that the interaction is mediated through this domain (Extended Data Fig 5g). Structural modeling of these candidate peptides onto the MAGE-A3 MHD suggests that all peptides occupy the same surface region—a hydrophobic cleft between the two winged-helix motifs that define the MHD (Fig 2b)^11,24,25^.

Sequence comparison revealed that these peptides share physicochemical properties rather than strict sequence conservation (Fig 2c). Modeling the BAP18 peptide within this cleft provided additional evidence for its fit into the shallow hydrophobic pocket on MAGE-A3 (Fig 2d). Thus, the amphipathic motif occupies a conserved hydrophobic cleft on the MAGE-A3 surface, similar to the previously described PCF11-MAGE-A11 interaction^24^, where a MAGE-binding region of PCF11 resides within a continuous helical segment as well.

To determine whether full-length BAP18 protein interacts with MAGE-A3 in a cellular context, we developed a NanoBRET assay^26^. Coexpression of BAP18 fused to HaloTag and MAGE-A3-NanoLuc produced a ∼18-fold increased signal compared to the soluble Halo-TAG control. Switching the HaloTAG between the N- and C-termini of BAP18 as well as swapping the donor and acceptor configurations yielded similar results, supporting a robust binary interaction in cells (Fig 2f and Extended Data Fig 5h).

After demonstrating that full length BAP18 interacts with MAGE-A3 in cells, we investigated if this interaction occurs through the interface identified through modelling and biochemistry, using *in vitro* pulldowns with peptide competition. Avi-tagged, biotinylated MAGE-A3 or biotinylated BSA (negative control), immobilized on beads, were incubated with full-length BAP18 in the absence or presence of a competitor peptide from the Y2H screen (TPT1 amino acids 120-140). Full-length BAP18 bound strongly to MAGE-A3 but not to BSA and the interaction was reduced in the presence of the competitor peptide but not a non-binding peptide (Extended Data Fig 5i, j). Therefore, BAP18 binds MAGE-A3 directly through the interface defined *in silico*.

A mutagenesis peptide array was used to assess if altering physicochemical properties of the substrate peptide modulates binding to MAGE-A3. Each residue within the core 15-mer of the BAP18 peptide was systematically substituted with all 20 canonical amino acids. The array was probed with full-length MAGE-A3 (Fig 2e). Introduction of polar – particularly negatively charged – residues within the core region (positions 50– 56 in the full-length sequence) markedly reduced MAGE-A3 binding. Arginine at position 51 appeared to be particularly important, since substitution with any other residue strongly reduced MAGE-A3 binding. Structural modeling of the peptide within the MAGE-A3 MHD suggests that this arginine lies adjacent to a negatively charged residue on MAGE-A3, potentially stabilizing the interaction through electrostatic complementarity (Fig 2d). The array binding pattern suggests moderate tolerance for amino acid substitutions that maintain the overall hydrophobic character of the MAGE-A3 substrate binding peptide. Consistently, selected BAP18 mutations decreasing hydrophobicity impaired the interaction with MAGE-A3 in cells (Fig 2f), suggesting a gradient of binding strength dependent on physicochemical properties of the hydrophobic interface.

### BAP18 degradation requires MAGE-A3 binding and polyubiquitination

To further dissect the molecular mechanisms underlying MAGE-A3-dependent BAP18 binding, polyubiquitination, and degradation, we developed a fluorescent protein degradation reporter system. We engineered DLD-1-iMAGE-A3 cells to express BAP18-meGFP and mCherry bicistronically (via P2A cleavage)^27^ (Fig 3a). Doxycycline treatment induced a shift of BAP18-meGFP relative to mCherry levels, consistent with post-translational regulation. The slopes of the linear regression models fitted to meGFP/mCherry fluorescence signals in control or doxycycline treatment were nearly identical, indicating that degradation of the BAP18-meGFP fusion occurs across all expression levels (Extended Data Fig 6a, b). Upon MAGE-A3 induction, BAP18-meGFP levels decreased to ∼25% of control, similar to the effect observed for endogenous BAP18 in DLD-1-iMAGE-A3 cells (Extended Data Fig 6c). To test whether interfering with substrate binding impacts protein degradation, we introduced mutations in the BAP18 reporter, which were reducing BAP18 binding to MAGE-A3 (Fig 2e, f; M49E, L50K and R51E). Introduction of the L50K mutation decreased the overall stability of the BAP18-meGFP fusion protein, while mCherry levels remained unaffected. All three mutations did not fully abrogate MAGE-A3-dependent degradation, but the slope of signal distribution was significantly increased for M49E and R51K, suggesting that these mutants are not efficiently degraded when expressed at higher levels, possibly due to the less efficient binding (Extended Data Fig 6d, e).

**Figure 3.**
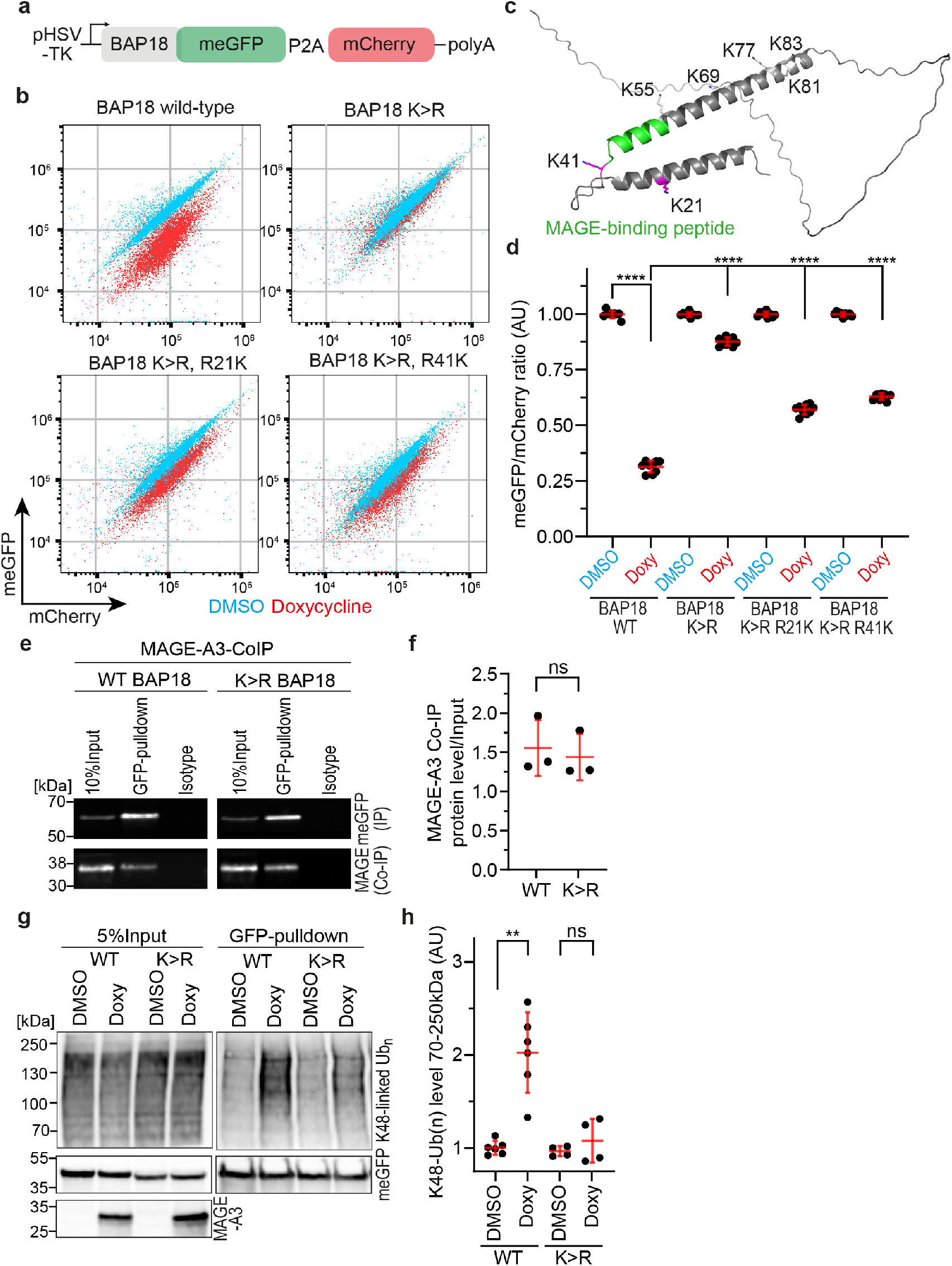
BAP18 degradation requires MAGE-A3 binding and lysine ubiquitination. **a**, Schematic of the bicistronic BAP18-degradation reporter. **b**, meGFP over mCherry flow cytometry fluorescence signals for the indicated versions of the BAP18 reporter expressed in DLD-1-iMAGE-A3 cells after 48-hour DMSO or doxycycline treatment. **c**, BAP18 protein model indicating all lysines in the structured part of the protein that were subject to mutagenesis. Lysines 21 and 41 are shown in purple. **d**, Quantification of BAP18-meGFP/mCherry ratios to assess BAP18 degradation as shown in panel b. Bars indicate mean ± s.d. Significance was tested using a two-tailed Mann-Whitney *U*-test (*P* = 4.11^-5^ (WT, doxy); *P* = 4.11^-5^ (K>R, doxy); *P* = 4.11^-5^ (K>R R21K, doxy); *P* = 4.11^-5^ (K>R R41K, doxy). **e**, Fluorescent immunoblot analysis of Co-IP of BAP18-meGFP with MAGE-A3 in DLD-1-iMAGE-A3 stably expressing WT or K>R reporter construct. **f**, Quantification of Co-IP as shown in panel e. Bars indicate mean ± s.d. Significance was tested using a two-tailed Mann-Whitney *U*-test (*P* = 0.40). **g**, Immunoblot analysis of K48-linked polyubiquitination of BAP18 WT and K>R reporter constructs by pulldown of the reporter in DLD-1-iMAGE-A3 cells after DMSO or doxycycline treatment. **h**, Densitometry quantification of K48-linked Ub(n) as shown in panel g, in a molecular weight range of 70-250 kDa. Significance was tested using a two-tailed Mann-Whitney *U*-test (*P* = 2.16^-3^ (WT); *P* = 0.99 (K>R)). Biological replicates: *n* = 9 (**b, d**), *n* = 3 (**e, f**), *n* = 6 (**g, h, WT**), *n* = 4 (**g, h, K**>**R**). Technical replicates: *n* = 1-2.

To identify mutations that fully disrupt degradation, we combined two mutations with a weak effect to synergistically impair the interaction between BAP18 and MAGE-A3 as interaction is mediated through an extended shallow surface on the MHD. The combination of L50E and A53R, selected to introduce polar amino acids in the otherwise hydrophobic stretch, abrogated MAGEA3-dependent degradation of BAP18-meGFP (Fig 2e and Extended Data Fig 6f, h). Conversely, increasing the hydrophobicity through the E46W-M49L double mutation (Fig 2e) slightly increased degradation compared to the wild-type (WT) sequence (Extended Data Fig 6g, h), suggesting that increased affinity facilitates degradation.

To identify the residues in the MAGE-A3 substrate binding cleft (SBC) that would impair BAP18 degradation by disrupting binding we selected mutations from other MAGE proteins previously identified to weaken substrate binding^24^ and inferred from MAGE-A3 MHD analysis. Comparison of the published MAGE-A3 MHD crystal structure (PDB: 4V0P)^28^ to an AlphaFold3 (AF3) model revealed close similarity (RMSD = 0.791; Extended Data Fig 7a). We therefore used AF3 modeling to assess the structural integrity of MHD mutants. Four mutations were introduced into the substrate binding cleft (SBC) of MAGE-A3, all exchanging hydrophobic to polar amino acids.

These included two mutations within the central region (F162R and V161R) and two adjacent to the cleft (L160R and L228R). AF3 modeling suggests near-identical folding of wild-type and mutant MHDs (Extended Data Fig 7b-f). Whereas transiently expressed wild-type MAGE-A3 downregulated BAP18-meGFP, all tested mutants showed reduced degradation activity (Extended Data Fig 7g-k). Mutations in the central portion of the SBC impaired degradation more strongly than those at the cleft periphery (Extended Data Fig 7l). These results indicate that the extended shallow hydrophobic cleft is required for substrate recognition and degradation.

UPS-mediated degradation requires polyubiquitylation at lysine residues. Therefore, we generated an all lysine (K) to arginine (R) mutant reporter construct, changing all lysines in the structured region of BAP18 (amino acids 1 to 91). The K>R reporter was not degraded upon MAGE-A3 induction. Reintroduction of either K21 or K41 was sufficient to restore ∼50% degradation. Both residues are located adjacent to the MAGE-A3 binding peptide (Fig 3b-d). Of note, other positions did not restore degradation (Fig 3c; Extended Data Fig 8a, b). While coimmunoprecipitation of MAGE-A3 with either wildtype or the K>R reporter showed no difference in interaction (Fig 3e, f), the K>R reporter lacked K48-linked polyubiquitination^29^ (Fig 3g, h), confirming that polyubiquitination of lysines is required for MAGE-A3-mediated degradation of BAP18.

### BAP18 loss induces cell migration

Having elucidated the molecular requirements of MAGE-A3-dependent BAP18 modulation, we set out to understand its biological relevance in cancer cells. For this, we selected A549 cells (lung adenocarcinoma, LU-AD), which display epithelial morphology, express low levels of MAGE-A3/6 (Extended Data Fig 1a, 3a, b), high levels of TRIM28 (Extended Data Fig 1b) and originate from a cancer type where MAGE-A3/6 reexpression is frequently observed^5^. Wild-type MAGE-A3 and MAGE-A3 V161R mutant (degradation-deficient) were expressed by stable transduction of parental cells (Extended Data Fig 7i, l, Extended Data Fig 9a, b) and WT MAGE-A3 but not the V161 mutant decreased BAP18 protein levels (Extended Data Fig 9a, b). Notably, WT MAGE-A3-overexpressing cells, unlike those with the V161R mutant, adopted a spindle-like shape (Extended Data Fig 9c, d), reminiscent of epithelial to mesenchymal transition (EMT)^30–32^.

Increased migratory capacity is a hallmark of EMT^33^. Live cell imaging revealed that MAGE-A3-overexpressing cells moved faster into a cell-free gap during a wound healing assay and frequently detached from the monolayer (Extended Data Fig 9c-e). Migrating cells showed an elongated shape with lamellipodia-like structures. Cells expressing the mutant MAGE-A3 V161R migrated less, mostly remaining in contact with the monolayer, similar to parental controls (Extended Data Fig 9c, d). Thus, MAGE-A3 expression in A549 cells promotes migration via its E3 ligase adaptor function.

Consistently, CRISPR/Cas9-mediated BAP18 KO in A549 cells (Extended Data Fig 9f, g) promoted a morphological change similar to MAGE-A3-overexpression. After BAP18 KO, A549 cells frequently exhibited an extended, spindle-like chape, accompanied by F-actin rich lamellipodia formation (visualized by stably expressed LifeAct-meGFP^34^), indicating a pronounced front-back polarity along the direction of migration (Fig 4a, b). BAP18 KO was also sufficient to promote cell migration in a wound healing assay (Fig 4c, d) and detachment of cells from the monolayer at the cell-free gap boundary (Fig 4e, Supplementary Movies 1 (wild-type), 2 (BAP18 KO)). The faster wound healing was not due to increased proliferation, as the growth rates of control and BAP18 KO cells were unchanged over 72 hours (Extended Data Fig 9h). To confirm the increased migratory behavior, we labeled control and BAP18 deficient cells with fluorescent histone 2B and tracked individual cells at the edge of a cell-free gap. Individual BAP18 deficient cells showed greater migratory potential, as indicated by higher cumulative displacement (Fig 4f, Extended Data Fig 9i). These findings suggest that BAP18 loss is a key mediator of MAGE-A3/6-induced effects in cancer and could support the metastatic phenotype linked to MAGE expression in tumors. The observed phenotypical transformation is accompanied by actin cytoskeleton remodeling, a critical indicator of cancer cell migration and transition toward a mesenchymal cell state^35,36^.

**Figure 4.**
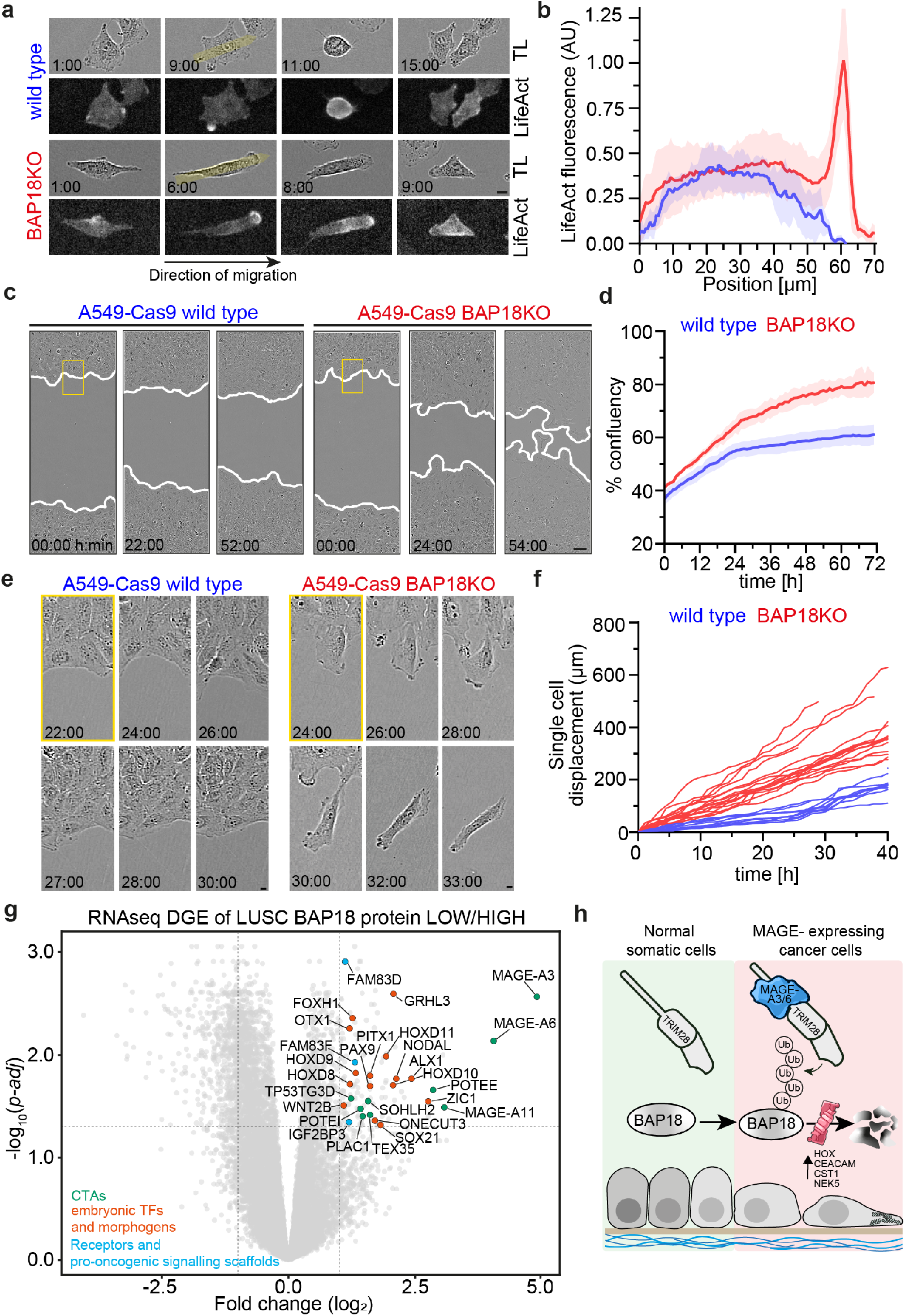
BAP18 downregulation induces cell migration. **a**, Representative images of A549 wild type or BAP18 KO cells expressing LifeAct-meGFP. Yellow shaded arrows indicate line profiles for LifeAct quantification. **b**, Quantification of LifeAct signal along the major axis of cells as indicated by shaded yellow arrows in a. Signal was normalized to the leading-edge signal of BAP18 KO cells. *n* = 12 cells (wild type) and *n* = 15 (BAP18 KO). **c**, Representative images of a wound-healing assay of parental or BAP18 KO A549 cells. Yellow rectangles indicate insets shown in panel e. **d**, Quantification of cell migration into the cell-free gap shown in panel c. Data are mean ± s.d. **e**, Insets of wild type or BAP18 KO A549 cell morphology at the border of the cell-free gap shown in panel c. **f**, Quantification of single cell cumulative displacement of A549 H2B-mCherry BAP18 KO cells compared to wild type. Data shows individual cell tracks. *n* = 10 cells (wild type) and *n* = 18 cells (BAP18 KO). **g**, Volcano plot displaying DEGs between bottom and top quartiles of BAP18 expression in LUSC tumor samples (30 tumors each). Significance and fc-thresholds are indicated as dashed gray lines (FDR adjusted *P*values < 0.05; |log2FC| > 1). Selected highly upregulated genes are labeled in green (CTAs), orange (embryonic TFs and morphogens), and light blue (receptors and pro-oncogenic signaling scaffolds). **h**, Model of MAGE-A3/6-TRIM28 dependent BAP18 degradation through the UPS. Loss of BAP18, induced through MAGE-A3/6, leads to a morphological change of cells and a migratory phenotype and weakens the epithelial cell adhesion. Biological replicates: *n* = 3 (**a-f**). Technical replicates: *n* = 1-3. Scale bars, 100 µm (**a, c**), 1 µm (**e, f**). Time is shown as h:min (**a**,**c**,**e**).

Since BAP18 is implicated in transcriptional regulation as part of the human NuRF/BPTF complex^37,38^, we performed RNA-seq of BAP18 KO A549 cells. Loss of BAP18 caused upregulation of 46 genes (adjusted *P*-value < 0.05, log_2_FC > 1), including several highly upregulated transcripts that are known to promote cell migration in various cancers (Extended Data Fig 9j). Among them, CST1^39^ and AQP3^40^ are reported to directly enhance migratory capacity, while NEK5, a kinase implicated in hemidesmosome destabilization^41^, may simultaneously weaken cell-cell adhesion. Two members of the carcinoembryonic antigen-related cell adhesion molecule (CEACAM) family, CEACAM 5 and 6, were also strongly upregulated. Both are cell surface proteins linked to metastasis^42^. Moderately upregulated genes included HOX cluster members (HOXD4 (fc=2.42, P = 2.15^-4^), HOXB2 (FC=2.23, P = 4.16^-4^)) and ID4 (FC=1.28, P = 3.46^-5^), a dominant-negative bHLH transcription factor capable of suppressing differentiation-associated programs^43^. These changes may indicate activation of developmental pathways and partial de-differentiation. The fact that MAGE-A genes, including A3 and A6, were reporter to be expressed in mesenchymal stem cells^44^, but not differentiated somatic tissue, suggests that BAP18 loss might be the functional link to how MAGE-A3/6 exert their function during normal organismal development and oncogenesis.

We observed a similar pro-migratory phenotype and morphological changes upon BAP18 KO in two independent MAGE-negative cells, FaDu (head and neck squamous cell carcinoma, HNSCC) and DLD-1 (Extended Data Fig 10a-l), suggesting that the role of BAP18 might be shared across various cancer types.

To study the translational relevance of our findings, we queried the proteogenomic LUSC dataset^21^. Differential gene expression analysis between the lowest and highest 25% of BAP18-expressing tumors identified 137 upregulated genes (adjusted *P*-value < 0.05, |log_2_FC| > 1; Fig 4g). MAGE-A3 and MAGE-A6 were the top differentially expressed genes, confirming the inverse correlation with BAP18. Thus, BAP18 protein level-mediated tumor stratification accurately reflects MAGE-A3/6 expression. Among the DEGs, a group of genes was linked to squamous cell identity. Notably, the transcription factor GRHL3, reported to support a reversible migratory state in progenitor epidermal cells^45^ and to induce migration in epithelial tumors^46^, was upregulated in BAP18-low tumors as well as in BAP18 KO A549 cells (Fig 4g, Extended Data Fig 9j) in accordance with the observed increased migratory capacity of BAP18 KO cells.

Among the upregulated genes, we observed other CTAs, including MAGE-A11 and POTE family members, and several developmental and embryonic transcription factors (TFs) and morphogens, including 4 members of the HOX cluster (HOXD8, HOXD9, HOXD10, HOXD11) and the embryonic morphogens NODAL and WNT2B. This indicates that germline and developmental programs are broadly reexpressed in these tumors. The reemergence of HOX genes in cancers has been recognized across many cancer types including squamous cell carcinoma (SCC) to be a driver of cellular plasticity, inducing de-differentiation and aiding cancers in the acquisition and maintenance of stem-like and invasive behaviour^47,48^. In particular, HOXD10 and D11 were shown to promote invasion in head and neck squamous cancer^49^ and NODAL is known to be involved in embryonic development and tumor progression^50^. Taken together, our data suggests that MAGE-A3/6-induced BAP18 degradation promotes de-differentiation and plasticity, facilitating migratory behavior (Fig 4i).

## Discussion

CTA expression in cancer often occurs concomitantly, owing to their overlapping reactivation mechanism and genomic proximity, as exemplified by the MAGE cluster on the X-chromosome. Therefore, delineating the role of individual CTAs is often complicated by their overlapping expression patterns.

In this study, we investigate the molecular function of an prototypical CTA and describe BAP18 as a previously unrecognized, bona fide substrate of MAGE-A3/6, targeted for proteasome-dependent degradation. We directly link its downregulation to induction of a promigratory cellular phenotype. BAP18 degradation by MAGE-A3/6 is highly specific, as only a handful of additional proteins are significantly downregulated upon MAGE-A3 expression. Effective degradation requires not only binding but also the presence of lysines optimally positioned relative to the interaction interface. In contrast to more promiscuous E3 adaptors such as CRBN, our data indicate that the MAGE-A3/6–TRIM28 complex targets a narrow substrate set. Previously proposed substrates of the MAGE-A3/6-TRIM28 complex were not confirmed in our system, including FBP1^15^. We conclude that previously reported substrates are either regulated in a context-dependent manner or through secondary effects.

To characterize the function of MAGE-A proteins in cancer progression, an understanding of substrate specificity is essential. The high similarity across the MHD of MAGE-A proteins complicates mapping the substrate space. We show that single residue substitutions in MAGE-A3 SBC are sufficient to modulate BAP18 engagement and degradation, suggesting that minor changes within the substrate-binding interface shift substrate specificity. Alterations in proximity of the SBC in otherwise highly similar MHDs lead to distinct substrate interactions for MAGE-A11 (73% MHD similarity to A3, binding to PCF11) and MAGE-A4 (78% MHD similarity to A3, binding to RAD18). Our study provides a foundation for systematic mapping of distinct and overlapping binding partners across a broader set of MAGE proteins.

Expression of CTAs is frequently linked to immune evasive tumor states^19,20^, possibly reflecting the selective pressure imposed by immune surveillance. Our findings however indicate that MAGE-A3/6, beyond its immunological implications, exerts a direct, cell-intrinsic role in driving tumor aggressiveness by hijacking developmental-like programs. Analyses of CCLE and patient-derived proteogenomic datasets support MAGE-A3/6-dependent regulation of BAP18 across cancers. Overall, our results clarify the molecular basis of MAGE-A3/6 function as an oncogenic E3 ligase adaptor. Further mechanistic studies will be necessary to fully elucidate the transcriptional rewiring in response to MAGE-A3/6 reexpression during tumorigenesis.

## Acknowledgements and funding

The authors thank the staff and laboratory scientists at Boehringer Ingelheim RCV/Cancer Cell Targeting (CCT) Department for technical support. Proteomics analyses were performed by the Proteomics Facility at IMP/IMBA/ GMI using the VBCF instrument pool. We thank Elisabeth Roitinger and the entire VBCF Proteomics facility for their technical support and experimental design advice. We thank Peggy Stolt-Bergner and Patrick Werni for recombinant protein production and purification. We thank colleagues from Experimental Medicine and Computational Innovation Departments (Onur Kaya, Donat Alpar and Eirini Christodoulaki) for support with RNA sequencing and data analysis. We thank Gintautas Vainorius and Christian Umkehrer for useful technical and conceptual discussions. We thank Anna Obenauf and Robyn Leigh Schenk for comments on the manuscript.

This work has received funding from Boehringer Ingelheim and the Austrian Promotion Agency/ Österreichische Forschungsförderungsgesellschaft FFG (grant number 911784). The Structural Genomics Consortium is a registered charity (no. 1097737) that receives funds from Bayer AG, Boehringer Ingelheim, Bristol Myers Squibb, Genentech, Genome Canada through Ontario Genomics Institute [OGI-196], EU/ EFPIA/OICR/ McGill/KTH/Diamond Innovative Medicines Initiative 2 Joint Undertaking [EUbOPEN grant 875510], Janssen, Merck KGaA (aka EMD in Canada and US), Pfizer, and Takeda. DBL is supported by Natural Sciences and Engineering Research Council (NSERC) and the Canadian Institutes of Health Research (CIHR).

## Author contributions

M.W.G.S. and P.M. conceived the project, with input and conceptual advice from M.K. and R.A.N. M.W.G.S. ideated, designed and conducted all experiments, except for the Y2H-screen (M.K.), NanoBRET (P.B., D.B.-L.), Peptide array (M.W.G.S. with M.M.), time course and washout experiments, cellular degradation reporter assays, Co-IPs and polyubiquitination assays (M.W.G.S. with M.S.P.), mass spectrometry sample preparations (M.W.G.S. with R.W.K.) and computational analyses concerning regression and correlation analysis in cell lines and tumor samples (M.W.G.S. with N.B.). M.W.G.S., R.A.N. and P.M. prepared the manuscript, with input from all authors.

## Methods

### Cell lines, cell culture and drug treatments

All cell lines used in this study were tested regularly for mycoplasm contamination and validated by next-generation sequencing (NGS) and short tandem repeat profiling (STR). Master cell banks were generated and maintained by the central tissue culture (CTC) service at Boehringer Ingelheim RCV and fresh cell aliquots were used for each experiment and for each cell line engineering project. Cell lines were maintained using the following media: DLD-1, A-427, Colo 320DM, and NCI-H358 in RPMI (Pan Biotech™ P04-18047, or ATCC-formulated RPMI, Thermo Scientific Gibco™ #A10491); A-375, and OUMS-23 in DMEM (Sigma®, #D6429); LentiX HEK293 and HEK293T in DMEM supplemented with 1% (v/v) GlutaMAX (Thermo Scientific Gibco #35050061) and 1% (v/v) sodium pyruvate (Thermo Scientific Gibco™, 11360039); A549 and HeLa S3 in F-12K Nutrient Mix (Thermo Scientific Gibco™, 21127-022); BT-20, Hep G2, RKO, SK-CO-1, SK-MES-1, and FaDu in EMEM (Sigma®, #M4655) supplemented with 1% (v/v) GlutaMAX and 1% (v/v) sodium pyruvate (and additionally 1% (v/v) MEM Non-essential amino acids for BT-20); and HCT 116 in McCoy’s 5A (Modified, Thermo Scientific Gibco™, #16600082). Depending on the expression constructs, cell lines were maintained in media supplemented with the following antibiotic combinations: DLD-1-RIEH-iMAGE-A3, DLD-1-RIEH-iMAGE-A6 (400 µg/mL hygromycin B (Thermo Scientific Gibco™ #10687010), 2 µg/mL puromycin (Thermo Scientific Gibco™ #A1113803)); DLD-1-RIEH-iMAGE-A3-pHSV-TK-BAP18 reporters (400 µg/mL hygromycin B, 2 µg/mL puromycin, 6 µg/mL blasticidin S (Thermo Scientific Gibco™ #A1113903)); DLD-1-RIEH-iMAGE-A3-pEF1a-BAP18 reporters (400 µg/mL hygromycin B, 2 µg/mL puromycin, 10 µg/mL blasticidin S); DLD-1_pHSV-TK-BAP18 reporters (6 µg/mL blasticidin S); A549-RIEH-Cas9 (parental) (500 µg/mL hygromycin B, 1 µg/mL puromycin); A549-RIEH-Cas9-MAGE-A3 and A549-RIEH-Cas9-MAGE-A3(V161R) (500 µg/mL hygromycin B, 1 µg/mL puromycin, 10 µg/mL blasticidin S); A549-Cas9 and FaDu-Cas9 (1 µg/mL puromycin); A549-Cas9-dgNEG/dgBAP18 and FaDu-Cas9-dgNEG/dgBAP18 (1 µg/mL puromycin, 500 µg/mL geneticin); DLD-1-Cas9 (2 µg/mL puromycin); DLD-1-Cas9-dgNEG/dgBAP18 (2 µg/mL puromycin, 500 µg/mL geneticin); A549-Cas9-dgNEG/dgBAP18-H2B-mCherry and A549-Cas9-dgNEG/dgBAP18-LifeAct-meGFP (1 µg/mL puromycin, 500 µg/mL geneticin, 10 µg/mL blasticidin S). Media were supplemented with either 10% Tet-system approved FCS (Takara, #631106) for experiments using doxycycline inducible cell lines or conventional FCS (Thermo Scientific Gibco™, #26140079) for all other experiments. All cells were cultured at 37°C in a 5% CO_2_ containing humidified atmosphere. For inducing expression of MAGE-A3 or A6 in doxycycline inducible expression DLD-1 cell lines, cells were treated with doxycycline hyclate solution at indicated concentrations (Sigma, #D9891). To inhibit the proteasome, cells were treated with 10 µM MG132 (Sigma, #M7449).

### Generation of inducible expression, genetically engineered and reporter cell lines

All plasmids used in this study were purchased from Genscript. Cell lines expressing fluorescent markers, constitutive or inducible expression cassettes were generated by stable integration using lentiviral transduction. Lentiviral particles (pantropic VSV-G or ecotropic pseudotyped) were generated using using Lenti-X™ one-step lentivirus packaging system single shots (Takara, #631275 or #631278), using Lenti-X™ HEK293T cells (Takara, #632180) as virus producing cells, according to manufacturer’s guidelines. Transduction of target cells was performed in the presence of 8 µg/mL polybrene infection reagent (Sigma™ TR-1003-G). Cell pools were selected using the appropriate antibiotic, or optionally subcloned by limiting dilutions in 96-well plates to obtain single cell clones. Generation of ecotropic pseudotyped lentiviral recipient cell lines (RIEP/RIEN/ RIEH) was conducted as described previously^51,52^. TRIM28 KO cell lines were generated by transient overexpression of all-in-one Cas9-TagBFP2 dual gRNA plasmid, containing a dual hU6-mU6 guideRNA expression cassette for increased on-target effect^53^ (TRIM28 sgRNA1: CGCCATTCACACGCCGCCGG, TRIM28 sgRNA2: AGAGCGCCTGCGACCCGAGA). 48 hours after transfection, cells were single-cell sorted on a CytoFLEX SRT benchtop cell sorter (Beckman Coulter Life Sciences), equipped with a 405 nm violet laser and a 450/45 nm bandpass filter for detecting TagBFP2 fluorescence, using the instrument’s cyclone movement system to deposit TagBFP2 positive cells into 96-well plates. Loss of TRIM28 expression in isolated clones was verified by simple western capillary electrophoresis immunoassays and functionally assessed by rescue of BAP18 degradation.

### Immunoblotting

Protein samples for immunoblot analysis were collected either by lysis of cells directly in 6-well multi-well cell culture plates or by harvesting cell pellets in 1.5 mL microcentrifuge tubes. Cells were lysed in 1X RIPA lysis and extraction buffer (Thermo Scientific Pierce™ #89901), supplemented with Roche cOmplete™ Mini protease inhibitor-Cocktail (Merck/Roche, #11836153001) and 1X HALT Protease/Phosphatase inhibitor cocktail (Thermo Scientific™, #87786), lysates clarified by centrifugation at 18.000 x g at 4°C for 15 minutes and protein concentration in the supernatant determined by BCA assay (Thermo Scientific Pierce™ BCA Protein Assay Kit, #23225). Colorimetric BCA reaction was quantified against a BSA standard (Thermo Scientific Pierce™ bovine serum albumin standard, ampules, 2 mg/mL, #23209) by measurement on a Perkin Elmer VICTOR X Light plate reader. Protein samples were separated using 4-12% Criterion™ XT Bis-Tris precast gels, 18-well, with 1X XT MES Running Buffer System (Bio-Rad, #1610789). Following electrophoresis, proteins were transferred onto a Trans-Blot Turbo 0.2 µm nitrocellulose membrane (Bio-Rad, #1704159), for chemiluminescence detection, or a Trans-Blot Turbo 0.45 µm LF PVDF membrane (Bio-Rad, #1704275), for fluorescent detection. Transfer was carried out using Bio-Rads turbo blot rapid transfer protocol, using the 7-minute, mixed MW transfer protocol. Nitrocellulose membranes were blocked in 5% (w/v) NFDM (Millipore™, 70166) in TBS-T with 0,05% Tween 20 (Bio-Rad, #1706435 and #1610781) and LF-PVDF membranes were blocked in Intercept™ (TBS) blocking buffer (LI-COR, #927-60001) for 3 hours at room temperature. Membranes were cut into appropriate molecular weight range slices and incubated with primary antibodies over night at 4°C, using 5 % (w/v) NFDM in TBS-T for chemiluminescence detection and Intercept® (TBS) antibody dilution buffer (LI-COR, #927-65001). For detection of K48-linked poylubiquititin, 3 % BSA (Sigma®, #A7030) solution in TBS-T was used as blocking and antibody dilution buffer.

Following antibodies were used: Vinculin (EPR8185, Abcam, ab129002, rabbit recombinant, 1:2500) MAGE-A3/6 (Cell Signaling Technology, CS25800, rabbit polyclonal, 1:2000), BAP18 (Bethyl Laboratories, A304-207A, rabbit polyclonal, 1:1000), GFP (proteintech/ chromotek, #pabg1, 1:2500), K48-linkage specific polyubiqitin (Cell Signalling, #8081, 1:1000). Membranes were washed 5 times with TBS-T and incubated with secondary antibodies, either goat anti-rabbit IgG-HRP conjugate (Bio-Rad, #1706515, 1:3000) or IRDye® 800CW goat anti-rabbit IgG (LI-COR, #926-32211, 1:20.000) for 1.5 hours at room temperature, followed by three washes with TBS-T. Enhanced chemiluminescence signal was developed with Western Lightning Plus Chemiluminescence substrate (Revvity, #NEL10300) and detected on a ChemiDoc MP (Bio-Rad). Near-infrared secondary antibodies were detected on a LI-COR Odyssey CLx near-infrared imaging system, using auto-settings for laser intensity, 169 µm resolution, high quality and 0.00 mm focus offset (for membrane detection format). Images were quantified using ImageStudio version 5.5 (LI-COR), using a top/bottom media background correction for detected bands, with a border width of 5.

### Simple western capillary immunoassay

For measurement of DLD-1-iMAGE-A3/6-time courses and washout experiment, as well as for measurements in TRIM28 KO DLD-1-iMAGE-A3 cells and MG132 proteasome inhibition treatments, cells were plated in 6-well dishes, at 150k cells per well in 2 ml medium. For time course/washout experiments, cells were treated with doxycycline for the indicated durations. Treated cells were washed once with PBS (Thermo Fisher Gibco™, #14190144) and lysed in 1X RIPA lysis and extraction buffer (Thermo Scientific Pierce™, #89901) supplemented with Roche cOmplete™ Mini protease inhibitor-Cocktail (Merck/Roche, #11836153001) and 1X HALT Protease/Phosphatase inhibitor cocktail (Thermo Scientific™, #87786). Protein lysates were clarified by centrifugation at 18.000 x g at 4°C for 15 minutes and protein concentration in the supernatant determined by BCA assay (Thermo Scientific Pierce™, BCA Protein Assay Kit, #23225). Colorimetric BCA reaction was quantified against a BSA standard (Thermo Scientific Pierce™ bovine serum albumin standard, ampules, 2 mg/mL, #23209) by measurement on a Perkin Elmer VICTOR X Light plate reader.

WCE protein samples were analyzed on a Jess™ Simple Western system (Protein Simple, Bio-Techne) suing the 12-230 kDa Separation Module (Portein Simple, Bio-Techne, #SM-W001). Samples were prepared following manufacturers guidelines, using 5 µg WCE per capillary, and denatured for 5 minutes at 98°C. The following antibodies were used, as solution in Antibody Dilution Buffer 2: MAGE-A3/6 (Cell Signaling Technology, CS25800, rabbit polyclonal, 1:50), BAP18 (Bethyl Laboratories, A304-207A, rabbit polyclonal, 1:50), TRIM28/Kap1 (Novus Biologicals, NBP500-158, rabbit polyclonal, 1:50). For total protein normalization, assays were run using the total protein normalization (TPN) normalization assay module for Jess (ProteinSimple, Bio-Techne, #AM-PN01). This module enables quantification of total protein signal within each capillary for normalizing the antibody staining derived chemiluminescence signal against. Electropherogram data analysis was performed using Compass Software (ProteinSimple, version 6.1.0.).

### In-cell western plate-based immunofluorescence

Cells were seeded in 96-well black wall/clear bottom TC plates with poly-D-lysine coated surface (Thermo Scientific™, #152037) at 10.000 cells/ well in 100 µl medium and allowed to adhere for 8 hours prior to doxycycline addition. Doxycycline dose titrations were performed by dispensing doxycycline hyclate solution (1 mg/mL in DMSO, Sigma D9891) directly onto plated cells in 96-well plates using a Tecan D300e digital dispenser. Following doxycycline treatment for 48 hours, cells were washed twice with room temperature PBS (Thermo Fisher Gibco™, #14190144) and fixed with 4% formaldehyde solution (Thermo Scientific Pierce™, #28906) in PBS for 10 minutes at room temperature. Afterwards, cells were treated with quenching solution (25 mM TRIS-HCl pH 7.4 in PBS) for 15 minutes, followed by permeabilization with 0.2% Triton X100 (Sigma, #93443) for 10 minutes at room temperature. Cells were then incubated in blocking buffer (0.45 µm filtrated 2% BSA in PBS) for 2 hours followed by incubation with primary antibody (anti-MAGE-A3 [EPR19065], Abcam, ab223162, rabbit recombinant, 1:500 or anti-BAP18/C17orf49, Invitrogen, #PA5-31990, 1:1000) in antibody-dilution buffer (0,45 µm filtrated 2% BSA solution in PBS-T containing 0,1% Tween 20 (Sigma®, #P1379) overnight at 4°C. Plates were washed 5 times with PBS-T, and incubated with secondary antibody (LI-COR IRDye® 800CW goat antirabbit IgG, 926-32211, 1:800) together with CellTag™ 700 stain (LI-COR, 926-41090, 1:500) to normalize to and account for well-to-well variation in cell number, in antibody dilution buffer. Plates were incubated for 2 hours at room temperatre, washed 3 times with PBS-T, once with PBS and placed upside down on a lint-free paper towel for 15 minutes to drain, protected from light. Plates were then sealed with transparent film and developed on a LI-COR Odyssey CLx near-infrared imaging system, using auto-settings for laser intensity, 169 µm resolution, high quality and 4.00 mm focus offset (for 96-well plate format).

### Plasmid transfection

For transient overexpression of MAGE-A3 proteins in DLD-1 cells stably expressing a fluorescent BAP18-meGFP/mCherry reporter, cells were plated in 6-well dishes at 100.000 to 200.000 cells/well in 2 ml medium. Cells were allowed to adhere overnight prior to transfection using jetOPTIMUS® DNA transfection reagent (Polyplus-Sartorius, 1010000025). Transfection was performed using the manufacturers guidelines, using 2 µg DNA:3 µl jetOPTIMUS per well of a 6-well dish. Cells were incubated for 48 hours prior to analysis. For generation of TRIM28 KO in DLD-1-iMAGE-A3 cells, plasmid transfections of all-in-one Cas9-P2A-TagBFP2 constructs containing a dual guideRNA expression cassette were performed using the Neon® transfection system (Thermo Scientific™) with the 100 µl tip format. Cells were grown to 70% confluency, harvested and washed with PBS and for each transfection 1×10^6^ cells were transfected with 5 µg plasmid in a suspension in 100 µL buffer R, using a voltage of 1300 V, pulse width of 20 ms and 2 pulses. Electroporated cells were dispensed into 6-well dishes containing fresh, selection antibiotic-free medium and were allowed to recover for 48 hours prior to further processing.

### Recombinant proteins

Recombinant MAGEA3 variants were expressed in *E. coli* BL21(DE3) or Rosetta2(DE3) cells transformed with the corresponding expression plasmids. For each construct, a 50 mL overnight preculture (grown in LB or 2×YT supplemented with the appropriate antibiotic) was used to inoculate 3 L of TB medium containing kanamycin or ampicillin. Cultures were grown at 37 °C until reaching an OD_600_ of approximately 1.1–1.9, induced with 0.25 mM IPTG, and subsequently incubated at 18– 20 °C overnight. Cells were harvested by centrifugation and stored at −20 °C until further processing. Cell pellets (63–74 g) were resuspended in lysis buffer (25–50 mM HEPES pH 7.5, 500 mM NaCl, 5% glycerol, 1 mM TCEP, and 10–20 mM imidazole) supplemented with protease inhibitor tablets and lysed by sonication on ice. Lysates were clarified by centrifugation (10 000 rpm, 45 min, 4 °C) and applied to Ni-NTA resin. After washing with lysis buffer, bound proteins were eluted using imidazole-containing buffer (typically 250 mM). Eluted fractions were subjected to overnight TEV protease digestion at 4 °C, followed by buffer exchange using a HiPrep Desalting 26/10 column. Cleaved samples were passed over Ni-NTA resin a second time to remove His-tagged contaminants and uncleaved protein, collecting the flowthrough or low-midazole fractions containing the target protein. Final purification was performed by size-exclusion chromatography on a HiLoad 16/600 Superdex 75 pg column. For the AVI-tagged MAGE-A3, invitro biotinylation was performed at ∼40 µM AVI-MAGE-A3 concentration using 1.2 µM GST-BirA ligase in the presence of 2.5 mM ATP, 5.5 mM MgCl□, and 175 µM biotin prior to the final size exclusion chromatography. Purified proteins were concentrated using Amicon 10 kDa MWCO centrifugal filters, analyzed by SDS-PAGE, and aliquots were snap-frozen and stored at −70 to −80 °C. Recombinant BAP18 was purchased from abcam (ab131691), split upon arrival into experiment sized aliquots, snap-frozen and stored at −70 to −80 °C.

### Yeast-2-hybrid screen

Yeast two-hybrid screening was performed by Hybrigenics Services, S.A.S., Evry, France (http://www.hybrigenics-services.com). The coding sequence of the bait protein fragment corresponding to human MAGE-A3 (amino acids 100-314, Uniprot ID P43357) was PCR-amplified and cloned into Hybrigenics bait vectors pB29, where the bait is fused to the N-terminus of LexA (resulting in N-MAGE-A3-LexA-C vector hgx6139v1). pB29 derives from the original pBTM116 vector^54,55^. The constructs were checked by sequencing and used as a bait to screen a random-primed human lung cancer derived cDNA library (“Human Lung Cancer_RP1”) constructed into pP6. The N-MAGE-A3-LexA-C bait construct was screened using a mating approach with YHGX13 (Y187 ade2-101::loxP-kanMX-loxP, mata) and L40DGal4 (matα) yeast strains as previously described^56^. 72.7×10^6^ interactions were analyzed in the presence of 200 mM 3-amino triazole (3-AT) to suppress bait autoactivation. The prey fragments of the positive clones were amplified by PCR and sequenced at their 5’ and 3’ junctions. The resulting sequences were used to identify the corresponding interacting proteins in the GenBank database (NCBI) using a fully automated procedure. A confidence score (PBS, for Predicted Biological Score) was attributed to each interaction as previously described^57^.

### Peptide array manufacturing and analysis

SPOT membranes^58,59^ were synthesized on an ResPep SL (Intavis Bioanalytical; now CEM) using the manufactures standard protocol including acetylation of the N-terminus to mimic intraprotein peptides. Peptide arrays were blocked and rehydrated in 3% BSA (Sigma A3070 in PBS-T (0.1% Tween-20) for 16 hours at 4 °C. Afterwards, arrays were incubated with recombinant fulllength MAGE-A3 at 1 µg/mL in blocking buffer for 2 hours at room temperature (RT). Arrays were then washed five times with PBS-T (0.1% Tween-20) and incubated with anti-MAGE-A3 antibody (ab223162, 1:2500 in blocking buffer). Following three washes with PBS-T, arrays were incubated with secondary antibody (800CW anti-rabbit IgG, LI-COR 926-32211, 1:2500; or HRP-anti-rabbit IgG, Bio-Rad #1706515, 1:3000) for 1 hour at RT. Finally, arrays were washed three times with PBS-T and developed either on an Odyssey CLx (Auto 800nm intensity, 169 µm resolution, high quality, 0.0 mm Offset) intensity or on a Bio-Rad ChemiDoc MP using Revvity ECL Plus Western Lightning (NEL103001EA). Arrays developed with fluorescent antibodies were quantified using ImageStudio version 5.5 (LI-COR).

### In vitro peptide and protein binding assays

Peptides with N-terminal Biotin-PEG(2x) were purchased from Genscript. Lyophilized peptides were reconstituted in DMSO at a concentration of 10 mg/mL, and snap-frozen in liquid N_2_ in experiment sized aliquots, which were stored at -70°C. For in vitro binding assays using peptides, peptides (BAP18: Biotin-{PEG2} PAGAKWTETEIEMLRAAVKR; TPT1: Biotin-{PEG2} TGAAEQIKHILANFKNYQFF; CCDC93: Biotin-{PEG2} AELIQYQKRFIELY-RQISAV; Non-binder: Biotin-{PEG2}STYRNVMEQFNPG-LRNLINL) were immobilized on streptavidin coupled magnetic beads (Thermo Scientific Pierce™, #88817). For each pulldown reaction, 15 µl magnetic bead slurry were incubated with 30 pmol biotinylated peptide in 300 µL binding buffer (PBS-T, 0,1% Tween 20, 0,1% BSA), rolling end-over-end in microcentrifuge tubes for 1 hour at room temperature. Beads were washed three times with binding buffer and resuspended in 300 µl ice-cold binding buffer containing 50 nM full-length MAGE-A3 or MAGE-A3 MHD (corresponding to residues 100-314). In vitro binding was performed for two hours at 4°C, rolling end-over-end. Beads were washed three times using IP lysis buffer as washing buffer (50 mM Tris-HCl pH 7.5, 150 mM NaCl, 1% NP-40, 1 mM EDTA, Thermo Scientific Pierce™, #87788). Bound protein was eluted by boiling for 10 minutes at 95°C using 2x XT sample buffer (Bio-Rad, #1610791), containing 100 mM DTT (Bio-Rad, #1610611). Eluate was analyzed using chemiluminescent immunoblot.

For in vitro binding assay of Avi-MAGE-A3 and full length BAP18 (Abcam, ab131691), equimolar amounts amounting to ∼0.057 nmol of 2 µg Avi-MAGE-A3 or 4 µg biotinylated BSA (Thermo Scientific Pierce™, #29130) were immobilized on 15 µl equibilibrated streptavidin couple beads (Thermo Scientific Pierce™, #88817) in 300 µl binding buffer (PBS-T, 0,1% Tween 20, 0,1% BSA) for 1 hour at 4°C rolling end-over-end in protein LoBind® microcentrifuge tubes (Eppendorf, #0030108116). Avi-MAGE-A3 or BSA coupled beads were washed three times with cold binding buffer and 5 µg BAP18 protein was added in 300 µl cold binding buffer.

For competition reactions, either DMSO, TPT1 peptide (TGAAEQIKHILANFKNYQFF) or non-binding peptide (STYRNVMEQFNPGL-RNLINL) were added to the binding reaction (each 100 µM), which was incubated at 4°C for 2 hours, rolling end-over-end. Beads were washed three times using IP lysis buffer as washing buffer (Thermo Scientific Pierce™, #87788). Bound protein was eluted by boiling for 10 minutes at 95°C using 2x XT sample buffer (Bio-Rad, #1610791), containing 100 mM DTT (Bio-Rad, #1610611). Eluate was analyzed using fluorescent immunoblot.

### NanoBRET assay

HEK293T cells were seeded at 4 × 10^5^ cells cells per 100 µL complete DMEM in 96-well white TC plates. Cells were incubated for 6 hours in a 37°C incubator prior to transfection. Instructions for the XtremeGene HP (Millipore Sigma, #6366236001) transfection system were followed, using 0.01 µg of the NanoLuc plasmid, 0.03 µg of the HaloTag plasmid, and 0.06 µg of empty pCDNA3.0 per well in optiMEM. For each NanoLuc construct, a corresponding control condition containing empty HaloTag vector was included. Conditions were all performed in duplicate wells. The following day, culture supernatants were removed and replaced with 40 µL of phenol red-free DMEM supplemented with 4% FBS and 1 µL mL^-1^ HT NanoBRET® 618 ligand (Promega #G9801). Two wells were used as no ligand controls where transfected cells were resuspended in phenol red-free DMEM supplemented with only 4% FBS. Cells were incubated for 30 minutes in a 37°C incubator in the presence of ligand. Nano-Glor substrate (Promega #N1572) was diluted in phenol red-free DMEM to 8 µL/ mL and 10 µL of the mix was added per well. Plates were read immediately (within 10 minutes) of adding the Nano-Glo® substrate on a CLARIOstar plate reader (BMGlabtech), utilizing mild orbital mixing (10 seconds) prior to scanning. Data for the 450 nm (donor) and 618 nm (acceptor) emissions were recorded and processed in accordance with the Promega technical manual (Literature no. TM616). Background signal from no lig- and controls was subtracted from all samples. Normalized BRET ratios were expressed as fold change over the empty HaloTag (“HT”) control for each NanoLuc construct (Fold over HT).

### Flow cytometry experiments and data analysis

Cells stably expressing fluorescent dual color reporters were detached with Accumax (Sigma, #A7089), washed once with PBS (Thermo Fisher Gibco™, #14190144) resuspended in FACS buffer (PBS supplemented with 2% FCS and 5 mM EDTA (Thermo Scientific Invitro-gen™, #15575020) and strained through a 40-µm mesh. Data were acquired on a CytoFLEX LX flow cytometer (Beckman Coulter) using the 488- and 561-nm laser lines to detect meGFP (BP525/40) and mCherry (BP610/20) fluorescence, with daily calibration performed using CytoFLEX Ready to Use Daily QC Fluorosphere beads (Beckman Coulter, #C65719). Enough live singlet events were recorded per sample to include at least 30.000 events in each quantificatio. Flow cytometry data analysis was performed using Becton Dickinson & Company (BD) FlowJo 10.10.0 with FloJoEngine v5.00000, Java Version 17.0.6+10-LTS, build number: 3b20e798. Gating strategies were defined a priori and applied uniformly across datasets. Dead cells, debris and doublets were excluded prior to quantification of meGFP and mCherry population medians.

Representative gating strategies for meGFP/mCherry reporter analysis and single-cell sorting (TRIM28 KO cell line generation) are provided in Supplementary Data Fig 1.

### Immunoprecipitation and Co-immunoprecipitation

For MAGE-A3 co-immunoprecipitation with BAP18, 5×10^6^ DLD-1-iMAGE-A3 cells stably expressing a BAP18-meGFP fusion protein (under EF1a promoter) were treated with 250 ng/mL doxycycline for 18 hours in a 10 cm dish. Cells were washed with PBS once and harvested by scraping. For immunoprecipitation, GFPtrap® magnetic agarose bead kits were used (proteintech/chromotek, #gtmak). The pulldown of meGFP protein was performed according to manufacturer’s guidelines. Briefly, cell pellets were lysed in 200 µl lysis buffer, supplemented with Roche cOmplete™ Mini protease inhibitor-Cocktail (Merck/Roche, #11836153001) and 1X HALT Protease/Phosphatase inhibitor cocktail (Thermo Scientific™, #87786) for 30 minutes on ice. Lysate was clarified by centrifugation at 18.000 x g for 15 minutes at 4°C and the supernatant mixed with 450 µl dilution buffer and split into two microcentrifuge tubes. Per pulldown, 25 µl GFP-trap® or V5-trap® (as isotype control) were added to the clear supernatant. Immunoprecipitation was performed for 45 minutes at 4°C, with rolling end over end. Bound protein was eluted by boiling for 10 minutes at 95°C using 2x XT sample buffer (Bio-Rad, #1610791), containing 100 mM DTT (Bio-Rad, #1610611). Eluate was analyzed using fluorescent immunoblot.

For detection of K48-linked polyubiquitination of BAP18, 5×10^6^ DLD-1-MAGE-A3 cells stably expressing BAP18 wild type or BAP18 K>R meGFP fusion proteins were treated with DMSO or 250 ng/mL doxycycline for 16 hours. For immunoprecipitation, GFP-trap® magnetic agarose bead kits were used (proteintech/chromotek, #gtmak). For detection of polyubiquitination, cells were collected using the same procedure, and pellets were lysed in 200 µl 1X RIPA buffer, supplemented with Roche cOmplete™ Mini protease inhibitor-Cocktail (Merck/Roche, #11836153001), 1X HALT Protease/ Phosphatase inhibitor cocktail (Thermo Scientific™, #87786), 10 mM iodoacetamide (Thermo Scientific™, #A39271) and 10 µM PR-619 (LifeSensors, #SI-9619). Cleared lysate was diluted with 450 µl dilution buffer and pulldown was performed with 50 µl equilibrated GFPtrap® slurry per pulldown sample, over night at 4°C with rolling end-over-end. Bound protein was eluted by boiling for 10 minutes at 95°C using 2x XT sample buffer (Bio-Rad, #1610791), containing 100 mM DTT (Bio-Rad, #1610611). Eluate was analyzed using chemiluminescent immunoblot.

### Wound healing migration assays and cell tracking

For assessing cell migration, the CytoSelect™ 24-well format wound healing assay was used (Cell Biolabs, #CBA-120-5). This assay utilizes proprietary inert inserts that generate a uniform 900 µm gap in each well. Cells were seeded into wells containing these inserts in a 24-well multi-well plate and allowed to adhere into a confluent monolayer around the insert overnight. For A549-RIEH-Cas9 parental, MAGE-A3 and MAGE-A3 (V161R), A549-Cas9-dgNEG/dgBAP18, DLD-1-Cas9-dgNEG/ dgBAP18, 450.000 cells; for FaDu-Cas9-dgNEG/ dgBAP18, 400.000 cell per well were seeded. Following removal of the insert, cells were washed twice with prewarmed medium and cell migration was monitored by time-lapse microscopy using an Incucyte S3 (Sartorius), imaging fields of view every hour at the cell gap boundary with 10x magnification. Incucyte S3 control and data collection was performed using Incucyte 2023A Rev2 software. Analysis of wound healing was perfomed using ImageJ Fiji (v1.53c, Java version 1.8.0_362, 64-bit), using a plugin for detecting confluency during gap closure^60^. Single cell tracking and track visualization was performed using TrackMate, available with ImageJ Fiji distribution^61^. LifeAct expressing cells were analyzed using the MorphoLibJ plugin^62^, which was used to derive aspect ratios and axis positions, along which LifeAct distribution was quantified with a line profile of 10 px width.

### Quantitative mass spectrometry by nanoLC-MS/MS and quantitative mass spectrometry data analysis

The nano HPLC system (Vanquish *Neo* UHPLC-System) was coupled to an Orbitrap Eclipse mass spectrometer, equipped with a FAIMS pro interface and a Nanospray Flex ion source (all parts Thermo Fisher Scientific). Peptides were loaded onto a trap column (PepMap Neo C18, 5 mm × 300 μm ID, 5 μm particles, 100 Å pore size, Thermo Fisher Scientific, #TFS-174500) using 0.1% TFA as mobile phase. After loading, the trap column was switched in line with the analytical column (Aurora Ultimate C18 25cm × 75 μm ID, 1.7 μm particles, 120 Å, operated at 50°C, Ionopticks, #25075C18). Peptides were eluted using a flow rate of 300nl/min (Aurora), starting with the mobile phases 98% A (0.1% formic acid in water) and 2% B (80% acetonitrile, 0.1% formic acid) and linearly increasing to 35% B over the next 180 min followed by an increase to 95% B in 1.7 min, a 4-min hold at 95% B, and re-equilibration with 2% B for three column volumes. The Eclipse was operated in data-dependent mode ‘Cycle Time’, performing a full scan (m/z range 380-1500, orbitrap resolution 120k, AGC target 4E5) at 4 different compensation voltages (CV -45, -55, -65, -75), followed each by MS/ MS scans of the most abundant ions for a cycle time of 0.75 sec per CV. MS/MS spectra were acquired using an isolation width of 1.2 m/z, AGC target of 1E4, intensity threshold of 1E4, maximum injection time of 35 ms, HCD with a collision energy of 30%, using the ion trap for detection in the rapid scan mode. Precursor ions selected for fragmentation (include charge state 2-6) were excluded for 20 s. The monoisotopic precursor selection filter and exclude isotopes feature were enabled.

For peptide identification, the RAW-files were loaded into Proteome Discoverer (version 2.5.0.400, Thermo Scientific). All MS/MS spectra were searched using MSAmanda v2.0.0.19924^63^. The peptide mass tolerance was set to ±10 ppm and fragment mass tolerance to ±400 mmu, the maximum number of missed cleavages was set to 2, using tryptic enzymatic specificity without proline restriction. Peptide and protein identification was performed in two steps. For an initial search the RAW-files were searched against the uniprot_reference_human_2024-05-31.fasta (20,518 sequences; 11,415,822 residues), supplemented with common contaminants and sequences of proteins of interest using carbamidomethylation of cysteine as a fixed modification. The result was filtered to 1 % FDR on protein level using the Percolator algorithm^64^ integrated in Proteome Discoverer. A sub-database of proteins identified in this search was generated for further processing. For the second search, the RAW-files were searched against the created sub-database using the same settings as above and considering the following additional variable modifications: oxidation on methionine, deamidation on asparagine and glutamine, glutamine to pyro-glutamate conversion at peptide N-terminal glutamine and acetylation on protein N-terminus. The localization of the post-translational modification sites within the peptides was performed with the tool ptmRS, based on the tool phosphoRS^65^. Identifications were filtered again to 1 % FDR on protein and PSM level, additionally an Amanda score cut-off of at least 70 was applied. Proteins were filtered to be identified by a minimum of 2 PSMs in at least 1 sample. Protein areas have been computed in IMP-apQuant^66^ based on the MaxLFQ algorithm^67^ considering unique and razor peptides. Resulting protein areas were normalized using iBAQ^68^ and sum normalization was applied for normalization between samples. Match-between-runs (MBR) was applied for peptides with high confident peak area that were identified by MS/MS spectra in at least one run. Statistical significance of differentially expressed proteins was determined using limma^69^. Visualization of qMS data as volcano plots was done using the web-based application VolcaNoseR^70^.

### Gene expression profiling (RNA-seq) and differential gene expression analysis

QuantSeq gene expression profiling and gene expression analysis were performed as previously described^71,72^. In short, biological triplicate cell samples of each condition were collected independently, and cell pellets snap frozen in liquid N_2_ and stored at -70°C. Total RNA samples were prepared using Qiazol lysis/ extraction reagent (Qiagen Cat no./ID 79306) followed by isolation using RNeasy Plus Universal Mini Kit with gDNA Eliminator treatment (Qiagen #73404) and RNA concentration and quality were determined using an Agilent Tapestation 4200. QuantSeq libraries were generated using the Lexogen QuantSeq 3′ mRNA-Seq V2 library prep kit FWD (Lexogen, #015.96) using 500 ng input RNA, according to the manufacturer’s instructions. Sequencing of multiplexed QuantSeq libraries was performed in-house on an Illumina NextSeq2000 (instrument ID: VH00399), generating single-end 76-bp reads with 12-bp i7 index reads. Raw reads were demultiplexed with the Illumina pipeline using 12-nt Lexogen UDI indice. Base calling and quality scoring were performed on-instrument using Illumina’s DRAGEN Bio-IT Platform, integrated into the NextSeq 2000 system software. Conversion of raw BCL files to zipped FASTQ files was carried out using DRAGEN BCL Convert (Illumina). Data processing was performed using a pipeline based on the ENCODE *Long RNA-seq* workflow, adapted for QuantSeq 3’ mRNA-Seq data. In brief, single-end reads were aligned to the *Homo sapiens* reference genome (hg38/GRCh38) using the STAR aligner (v2.5.2b), with soft-clipping enabled to remove residual adapter or poly(A)-derived sequences typical for QuantSeq libraries. Transcript annotation from Ensembl release 86 was used for quantification. Gene-level counts were obtained using featureCounts (v1.5.1). Quality control assessments were performed at the appropriate steps using FastQC (v0.11.5), picardmetrics (v0.2.4), and dupRadar (v1.2.2). Differential expression analysis was conducted using DESeq2 on the gene-level count matrix generated by featureCounts. Unless stated otherwise, significance was defined as an absolute log_2_fold-change ≥1 and a false discovery rate <0.05. Visualization of differentially expressed genes as volcano plots was performed using the web-based application VolcaNoseR^70^.

### CCLE cell line regression analysis

Quantitative proteomics data on 375 cell lines from the Cancer Cell Line Encyclopedia (CCLE), as published by Nusinow et al.^17^, was utilized for regression modeling of protein expression. Standard linear regressions were performed using R’s lm() function for every protein against MAGEA3/MAGEA6 (Uniprot IDs: P43357, 43360) protein expression. Proteins were filtered to retain those with expression data available in at least 100 cell lines and a sufficient overlap of at least 50 cell lines with MAGEA3/6 expression data. Multiple testing correction was applied using the fdrtool package and proteins with q-values < 0.05 were considered significantly associated.

### Differential gene expression analysis in LUSC tumor sets

For both proteomic and transcriptomic LUSC datasets^21^,108 tumor samples were analyzed. BAP18 protein abundance (gene C17orf49) was extracted from the proteomics dataset, and samples were ranked by expression; the top 30 expressers were assigned to the BAP18□high group and the bottom 30 to the BAP18□low group. Differential expression analysis was performed for matched RNA□seq data by subsetting each dataset to these 60 samples and constructing a HIGH vs. LOW matrix. Linear modeling and empirical Bayes moderation were carried out using the limma package (lmFit, eBayes)^69,73^. Significance cutoffs were set at |log□FC| > 1 and FDR < 0.05.

### Databases

RNAseq data for in-cell western panel cell lines was obtained from the web-based cancer genomics database Ordino^74^. CRISPR□based gene□dependency estimates and RNA expression data for CCLE cell lines were retrieved from the DepMap (https://depmap.org/portal) ‘Chronos’ and RNAseq datasets (Cancer Dependency Map, Broad Institute), using Chronos gene□effect scores from DepMap^75^ release 25Q4.

### Software version details

The Fiji integrated distribution/version of ImageJ used for analyses in this study was ImageJ v1.53c,, using Java v.1.8.0_362 (64-bit), with custom/published ImageJ plugins as indicated. Peptide helical wheel projection was generated using NetWheels peptide wheel web based tool ([NetWheels: Peptides Helical Wheel and Net projections maker](https://neutrophil-proteome.shinyapps.io/netwheels/))^76^. Volcano plot visualizations were generated using VolcaNoseR ([VolcaNoseR-Exploring volcano plots](https://huygens.science.uva.nl/VolcaNoseR/))^70^. For flow cytometry data analysis and figure panel layout, FlowJo™ v 10.10.0, using Java v 17.0.6+10-LTS build number 3b20e798 was used. For peptide sequence alignments, SnapGene v 8.0.3 was used. For analysis and visualization of protein structure files, The PyMOL Molecular Graphics System version v 3.1.0a0 was used (Open-Source Build). For depicting false-color surfaces with hydrophobicity, YRB color panel was used by installing the script in Pymol^77^. GraphPad PRISM 10.4.1 (627) was used for statistical testing and regression analysis except CCLE harmonized MS regression analysis.

### Statistical analysis and data reporting

No statistical methods were used to predetermine sample size. To test significance, the robust, nonparametric Mann-Whitney *U*-test was used, except the NanoBRET assays, for which a Dunnet’s multiple comparison test was used. When manual annotation was required, blinding precautions were made.

### Sample numbers and replication

Fig. 1a: qMS analysis of DLD-1-iMAGE-A3 cells (*n* = 3). Fig. 1 b, c: Representative example and quantification of fluorescent immunoblot analysis of Vinculin, MAGEA3 and BAP18 levels in DMSO (*n* = 6), 25 ng/mL doxycycline (*n* = 6) and 250 ng/mL doxycycline (*n* = 6) treated samples of DLD-1-iMAGE-A3 cells. Fig. 1d: Quantification of MAGE-A3 (*n* = 3) and BAP18 (*n* = 3) levels in 250 ng/mL doxycycline treated DLD-1-iMAGE-A3 cells over indicated durations. Fig. 1 e: Quantification of MAGE-A3 (*n* = 3) and BAP18 (*n* = 3) levels in 250 ng/ mL doxycycline treated DLD-1-iMAGE-A3 cells after washout for the indicated durations. Fig. 1f: Quantification of BAP18 protein levels (*n* = 6) after 250 ng/mL doxycycline treatment of DLD-1-iMAGE-A3 parental and TRIM28 KO cells. Fig. 1h: Harmonized MS and RNAseq levels of MAGE-A3/6 protein and BAP18 protein/RNA levels from *n* = 54 sampled tumors. Fig. 2e: Quantification of MAGE-A3 engagement with peptide of indicated composition for *n* = 2 peptide arrays. Fig. 2f: Quantification of NanoBRET ratio in cells expressing MAGE-A3-HaloTAG and BAP18-NanoLuc (NL) (*n* = 4), BAP18^M49E^-NL (*n* = 4), BAP18^L50K^-NL (*n* = 4) and BAP18^R51E^-NL (*n* = 4). Fig. 3b, d: Representative example scatterplot and quantification of flow cytometry analysis of BAP18-meGFP/mCherry wild type (*n* = 9), K>R (*n* = 9), K>R R21K (*n* = 9) or K>R R41K (*n* = 9) reporter constructs expressed in DLD-1-iMAGE-A3 cells after treatment with 250 ng/mL for 48 h. Fig. 3e, f: Representative fluorescent immunoblot and quantification of coimmunoprecipitation of MAGE-A3 with BAP18-meGFP wild-type (*n* = 6) or K>R (*n* = 6) reporter expressed in DLD-1-iMAGE-A3 cells. Fig. 3g, h: representative Immunoblot and quantification of K48-polyubiquitin levels of BAP18-meGFP wild-type (*n* = 6) and K>R (*n* = 4) reporter expressed in DLD-1-iMAGE-A3 cells. Fig. 4a, b: Representative example and quantification of LifeAct-meGFP distribution along the major axis of wild type (*n* = 12) and BAP18 KO (*n* = 15) A549 cells. Fig 4 c-e: Representative examples and quantification of wound healing of fields of view of wild type (*n* = 4) and BAP18 KO (*n* = 5) A549 cells. Fig. 4f: Quantification of single cell tracks of H2B-mCherry expressing wild type (*n* = 10) and BAP18 KO (*n* = 18) A549 cells. Fig. 4g: DEG analysis of LUSC tumors, comparing 30 BAP18 low and 30 BAP18 high (porein level) samples. Extended Data Fig. 1c: Representative example of in-cell western immunofluorescence analysis of MAGE-A3 expression (*n* = 9). Extended Data Fig. 1d: qMS analysis of DLD-1-iMAGE-A3 cells (*n* = 3). Extended Data Fig. 1e, f: Representative examples of simple western analysis of MAGE-A3 and BAP18 protein levels in DLD-1-iMAGE-A3 cells after doxycycline treatment or washout. Extended Data Fig. 2b: Representative examples of TRIM28, MAGE-A3 and BAP18 as well as total protein (TPN) levels in DLD-1-iMAGE-A3 wild type (*n* = 6) and TRIM28 KO cell (*n* = 6) samples. Extended Data Fig. 2 c, d: Representative example and analysis of Vinculin, MAGEA3 and BAP18 protein levels in DLD-1-iMAGE-A3 cells after DMSO (*n* = 4), doxycycline (*n* = 4) and doxycycline/ MG132 (*n* = 4) treatment. Extended Data Fig. 2 e, f: Representative example of in-cell western immunofluorescence analysis of MAGE-A6 expression (*n* = 9) and quantification of MAGE-3 and MAGE-A6 expression in DLD-1-iMAGE-A3 (*n* = 9) and DLD-1-iMAGE-A6 (*n* = 9) cells.

Extended Data Fig. 2g, h: Representative example and quantification of simple western analysis of MAGE-A6 (*n* = 3) and BAP18 (*n* = 3) levels in DLD-1-iMAGE-A6 cells. Extended Data Fig. 3a, b: Representative examples and quantification of MAGE-A3, BAP18 and CellTag700 levels (*n* = 8) in indicated cell lines. Extended Data Fig. 3d: Analysis of MAGE-A3/6 protein levels and BAP18 protein/RNA levels in *n* = 114 MMM tumors. Extended Data Fig. 4a: qMS analysis of DLD-1-iMAGE-A3 cells (*n* = 3). Extended Data Fig. 4b-i: Analysis of protein/RNA levels in LUSC (*n* = 54 tumor samples, except ALKBH2, *n* = 44 tumor samples) and MMM (*n* = 105 tumor samples, except p53, *n* = 86 tumor samples). Extended Data Fig. 5g: Representative examples of peptide pulldowns (*n* = 2). Extended Data Fig. 5h: Quantification of NanoBRET ratio in cells expressing MAGE-A3-NanoLuc and HaloTAG (*n* = 6), HaloTAG-BAP18 (*n* = 6) or BAP18-HaloTAG (*n* = 6). Extended Data Fig. 5i, j: Representative example and quantification of fluorescent immunoblot analysis of BAP18 binding to BSA (*n* = 6), Avi-MAGE-A3 (*n* = 6), Avi-MAGE-A3 in the presence of TPT1 peptide (*n* = 6) or non-binding peptide (*n* = 6). Extended Data Fig. 6a-c: Representative examples and quantification of slope (*n* = 9) and meGFP/mCherry ratios (*n* = 9) in DLD-1-iMAGE-A3 cells expression BAP18-meGFP/mCherry reporter. Extended Data Fig. 6d, e: Representative examples and quantification of meGFP/mCherry ratios in DLD-1-iMAGE-A3 cells expressing wild type (*n* = 9), M49E (*n* = 9), L50K (*n* = 9) or R51E (*n* = 9) BAP18-meGFP/mCherry reporter. Extended Data Fig. 6f-h: Representative examples and quantification of meGFP/mCherry ratios in DLD-1-iMAGE-A3 cells expressing L50E/A53R (*n* = 9) or E46W/M49L (*n* = 9) BAP18-meGFP/mCherry reporter. Extended Data Fig. g-l: Representative examples and quantification of DLD-1 cells expressing BAP18-meGFP/mCherry reporter, transiently expressing vector (*n* = 5), wild type MAGE-A3 *(n* = 5) or MAGE-A3 L160R *(n* = 5), V161R (*n* = 5), F162R (*n* = 5) or L228R (*n* = 5). Extended Data Fig. 8a, b: Representative examples and quantification of DLD-1-iMAGE-A3 cells expressing BAP18-meGFP/ mCherry K>R (*n* = 9), K>R R55K (*n* = 9), K>R R69K (*n* = 9), K>R R77K (*n* = 9), K>R R81K (*n* = 9) or K>R R83K (*n* = 9) reporters. Extended Data Fig. 9a, b: Representative example of vinculin, MAGE-A3 and BAP18 and quantification of fluorescent immunoblot analysis of BAP18 protein levels in parental (*n* = 4), MAGE-A3 (*n* = 4) or MAGE-A3 V161R (*n* = 4) expressing A549 cell samples. Extended Data Fig. 9c-e: Representative images and quantification of wound healing of fields of view of wild-type (*n* = 8), MAGE-A3 (*n* = 9) and MAGE-A3 V161R (*n* = 9) expressing A549 cells. Extended Data Fig. f, g: Representative example of vinculin and BAP18 and quantification of fluorescent immunoblot of BAP18 protein level in wild type (*n* = 6) and BAP18 KO (*n* = 6) A549 cell samples. Extended Data Fig. 9h: Quantification of CTG 2.0 proliferation of wild type (*n* = 8) and BAP18 KO (*n* = 8) A549 cells. Extended Data Fig. 9j: RNA seq DEG analysis of BAP18 KO (*n* = 3) over wild type (*n* = 3) A549 cells. Extended Data Fig. 10a, b: Representative example of vinculin and BAP18 and quantification of fluorescent immunoblot of BAP18 protein level in wild type (*n* = 6) and BAP18 KO (*n* = 6) FaDu cell samples. Extended Data Fig. 10c, d: Representative example of vinculin and BAP18 and quantification of fluorescent immunoblot of BAP18 protein level in wild type (*n* = 6) and BAP18 KO (*n* = 6) DLD-1 cell samples. Extended Data Fig. 10e-g, k: Representative images and quantification of wound healing of fields of view of wild-type (*n* = 4) and BAP18 KO (*n* = 6) FaDu cells. Ex-Extended Data Fig. 10h-j, l: Representative images and quantification of wound healing of fields of view of wild-type (*n* = 9) and BAP18 KO (*n* = 6) DLD-1 cells. All experiments in this study were performed in at least two biological replicates, and as often as indicated.

---

**Extended Data Figure 1.**
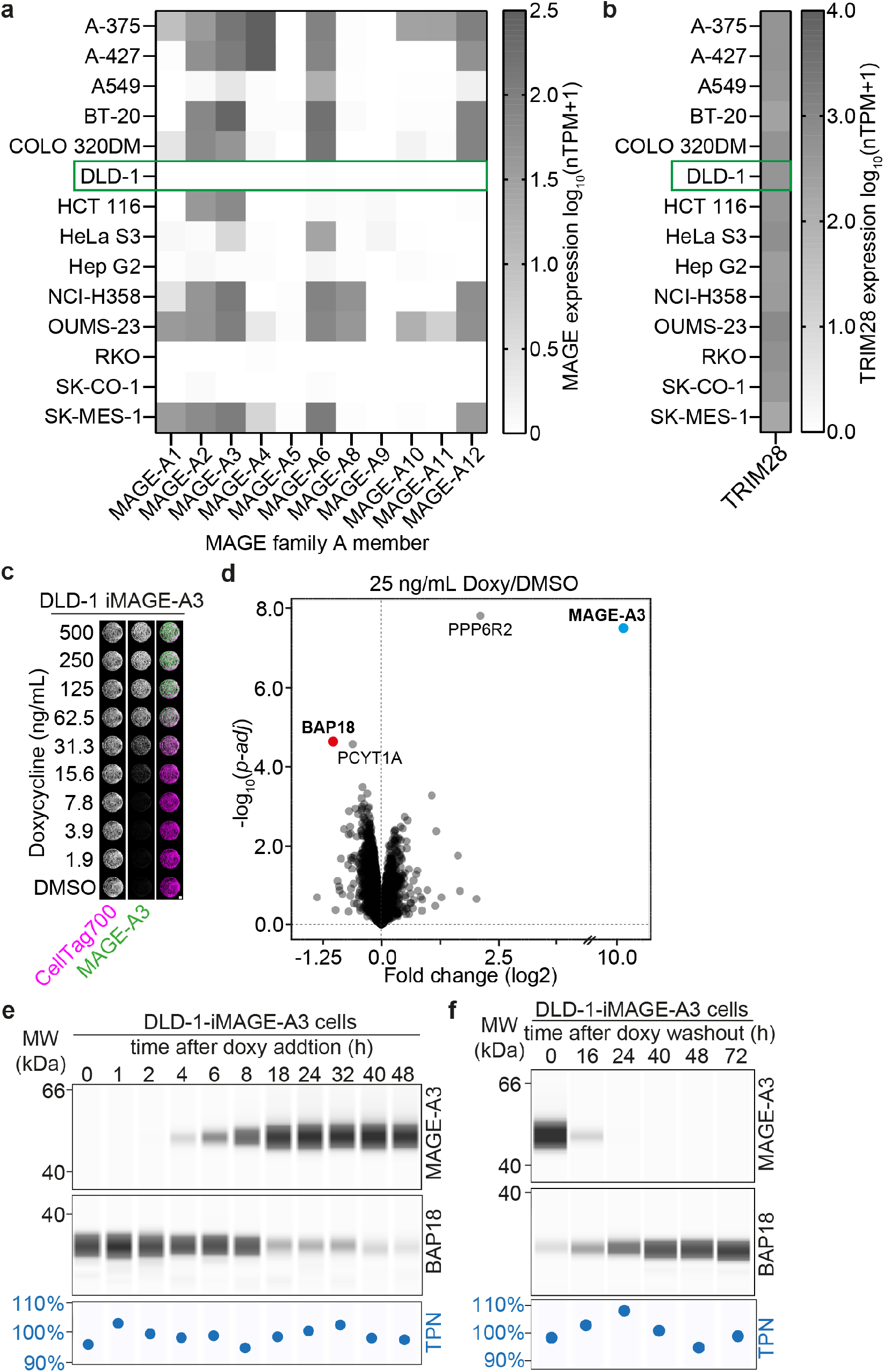
Generation and characterization of a model cell line for conditional MAGE-A3 over-expression. **a, b**, Heatmaps representing MAGE-A family and TRIM28 mRNA expression in a cell line panel. **c**, Immunofluorescence analysis of MAGE-A3 expression in DLD-1-iMAGE-A3 cells after treatment with varying doses of doxycycline for 48h. **d**, Volcano plot displaying qMS analysis of DLD-1-iMAGE-A3 cells after 40h treatment with 25 ng/mL doxycycline compared to DMSO. Top up- and downregulated protein indicated in bold. **e**, Simple western analysis of MAGE-A3, BAP18 and total protein (TPN) levels during time course after doxycycline addition as shown in Fig 1d. **f**, Simple western analysis of MAGE-A3, BAP18 and TPN levels after doxycycline washout as shown in Fig 1e. Biological replicates: *n* = 3 (**c, d**). Technical replicates: *n* = 1-3.

**Extended Data Figure 2.**
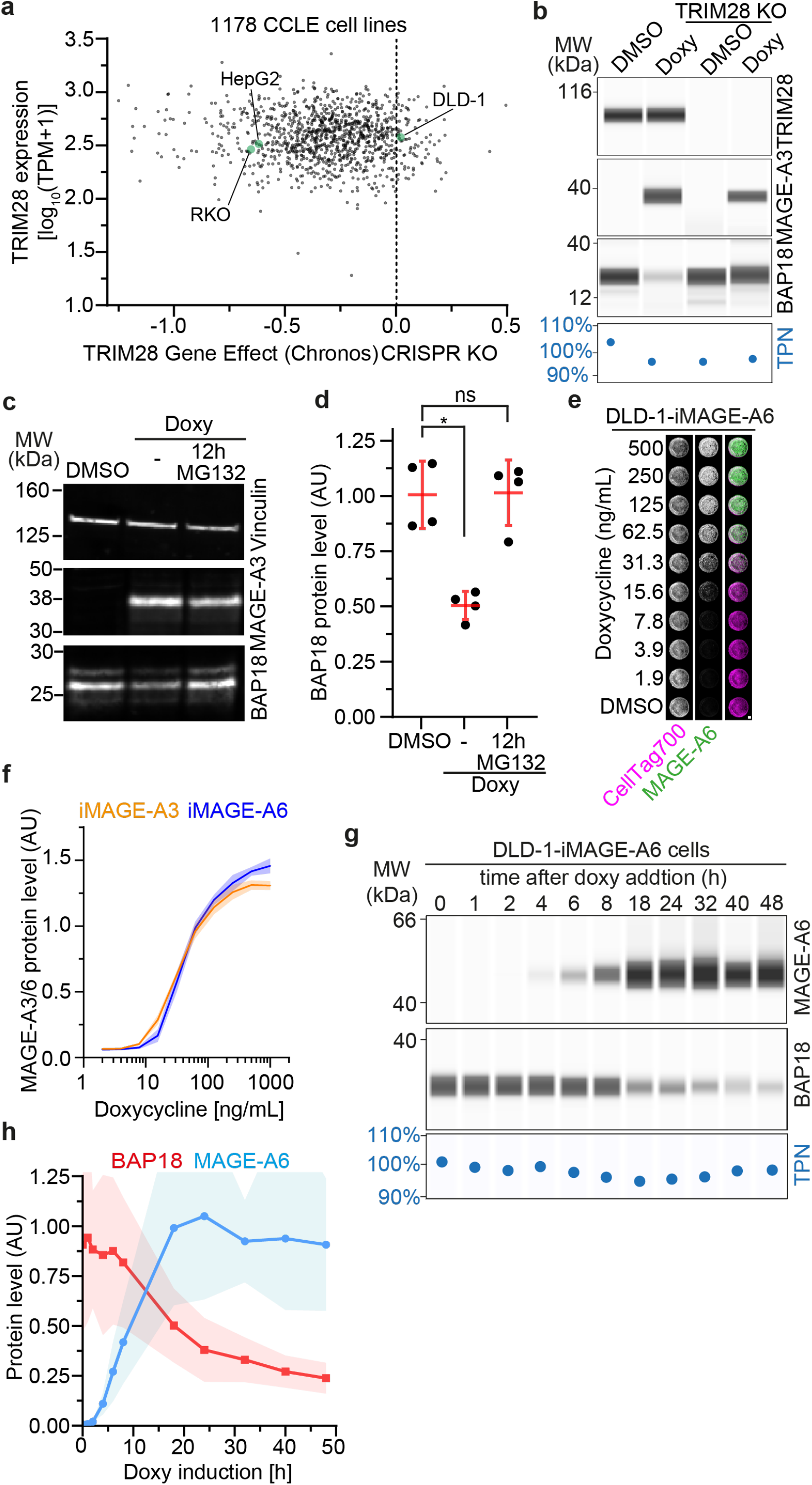
Investigation of TRIM28 and proteasome dependency of MAGE-A3 induced BAP18 degradation and analysis of paralog MAGE-A6. **a**, TRIM28 gene expression and TRIM28 KO (Chronos) score for 1178 CCLE cell lines. Data gathered by querying DepMap database in 25Q4. **b**, Simple western analysis of TRIM28, MAGE-A3, BAP18 and TPN levels in DLD-1-iMAGE-A3 and DLD-1-iMAGE-A3-TRIM28KO cells after DMSO or doxycycline treatment for 48 hours as shown in Fig 1f. **c**, Fluorescent immunoblot analysis of MAGE-A3 and BAP18 protein levels after DMSO, doxycycline or doxycycline and MG132 treatment. **d**, Quantification of fluorescent immunoblots shown in panel c. Bars indicate mean ± s.d. Significance was tested with a two-tailed Mann-Whitney *U*-test (doxy, *P* =2.86^-2^; doxy/MG132, *P* = 0.69). **e**, Immunofluorescence analysis of MAGE-A6 expression in DLD-1-iMAGE-A6 cells after treatment with varying doses of doxycycline for 48h. **f**, Quantification of MAGE-A3 and -A6 immunofluorescence shown in panel e and Extended Data Fig 1c. MAGE-A3 and -A6 levels are normalized to CellTag700. Data are mean ± s.d. **g**, Simple western analysis of MAGE-A6, BAP18 and total protein (TPN) levels during time course after doxycycline addition. **h**, Quantification of BAP18 and MAGE-A6 protein levels during time course. Data are mean ± s.d. Biological replicates: *n* = 6 (**b**), *n* = 4 (**c, d**), *n* = 3 (**e-h**). Technical replicates: *n* = 1-2.

**Extended Data Figure 3.**
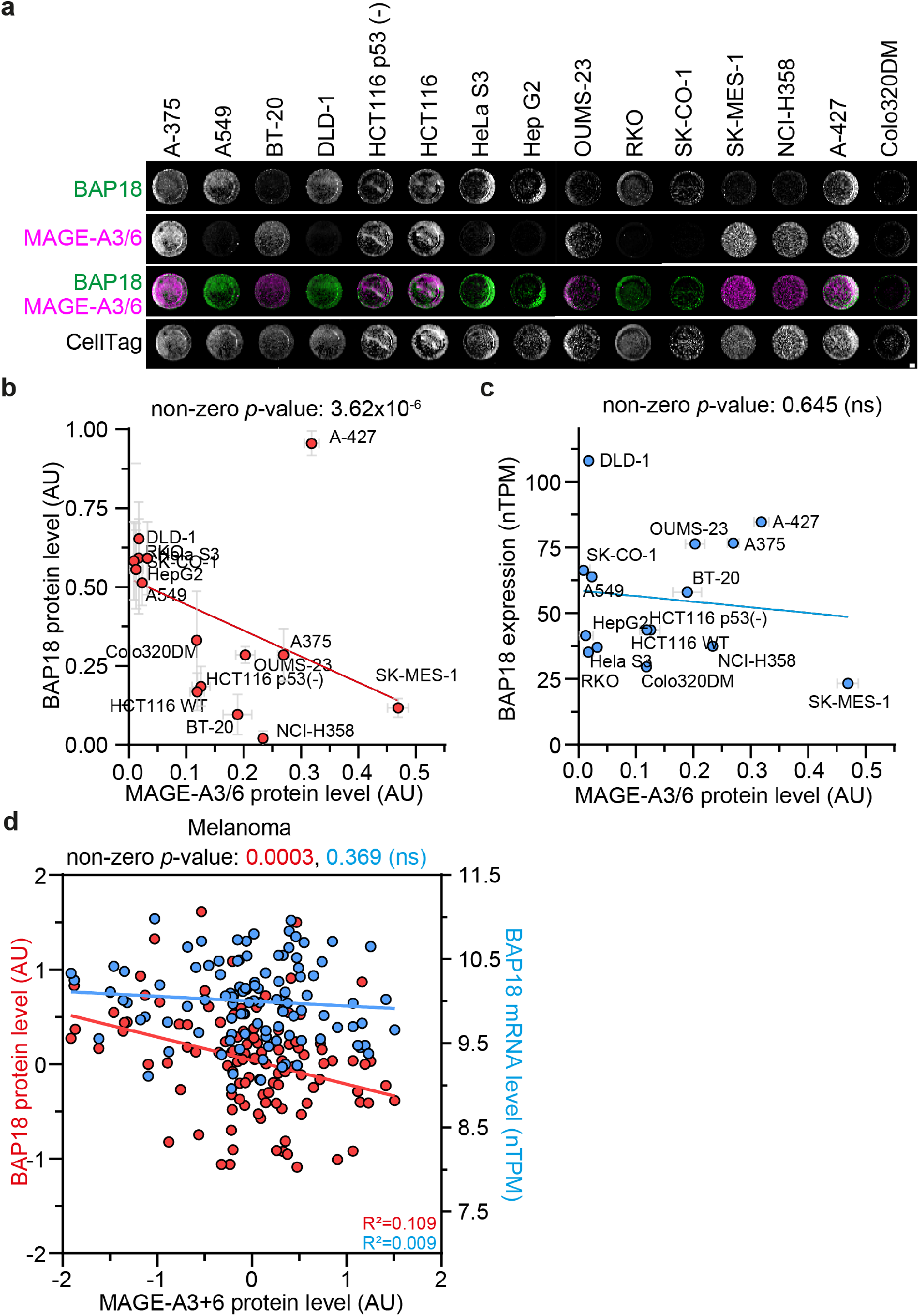
Correlation analysis of MAGE-A3/6 and BAP18 protein/RNA in cell line panel and metastatic malignant melanoma. **a**, Immunofluorescence analysis of MAGE-A3/6 and BAP18 protein levels in a panel of 15 cancer cell lines from varying indications. **b, c**, Regression analysis of BAP18 protein (**b**) or mRNA (**c**) against MAGE-A3/6 protein as shown in panel a (protein) or based on nTPM from RNAseq, Ordino database (mRNA). BAP18 and MAGE-A3/6 protein levels are normalized over CellTag700 counterstain. *P*-value indicates the deviation of linear regression line slope from zero. Protein data are mean ± s.d. **d**, Regression analysis of harmonized BAP18 protein and mRNA expression against MAGE-A3/6 harmonized protein expression in treatment naïve malignant melanoma tumor samples. *P-*values indicate significant/not-significant deviation of the linear regression line slope from zero. *n* = 115 tumors. Biological replicates: *n* = 2 (**a**-**c**). Technical replicates: *n* = 4.

**Extended Data Figure 4.**
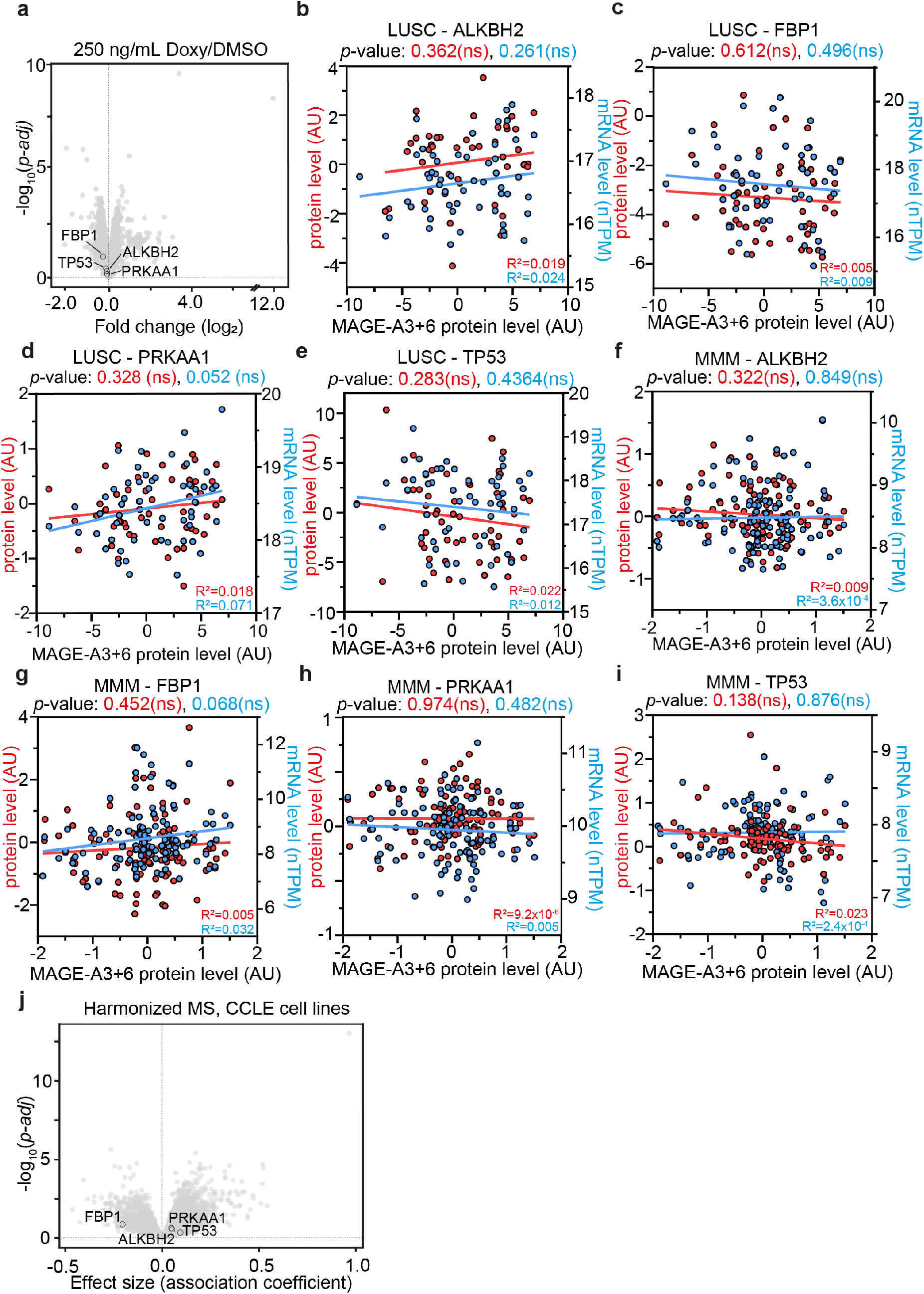
Correlation and regression analyses of other putative MAGE-A3/6 substrates. **a**, Volcano plot displaying qMS analysis of iMAGE-A3 DLD-1 cells after 40h treatment with 250 ng/mL doxycycline compared to DMSO. Putative other MAGE-A3 substrates are encircled. **b-e**, Regression analysis of harmonized protein and mRNA expression of the indicated proteins against MAGE-A3/6 harmonized protein expression in LUSC tumor samples. *P*-values indicate non-significant deviation of the linear regression line slopes from zero. *n* = 54 tumors, except for ALKBH2 protein: *n* = 44 tumors. **f-i**, Regression analysis of harmonized protein and mRNA expression of the indicated proteins against MAGE-A3/6 harmonized protein expression in treatment naïve metastatic malignant melanoma (MMM) tumor samples. *P-*values indicate non-significant deviation of the linear regression line slopes from zero. *n* = 115 tumors for protein analysis and *n* = 105 tumors for mRNA analysis, except for p53 protein *n* = 86 tumors. **j**, Regression analysis of every protein against MAGE-A3/6 expression in proteomics data from 375 CCLE cell lines. Putative other MAGE-A3 substrates are encircled. Biological replicates: *n* = 3 (a).

**Extended Data Figure 5.**
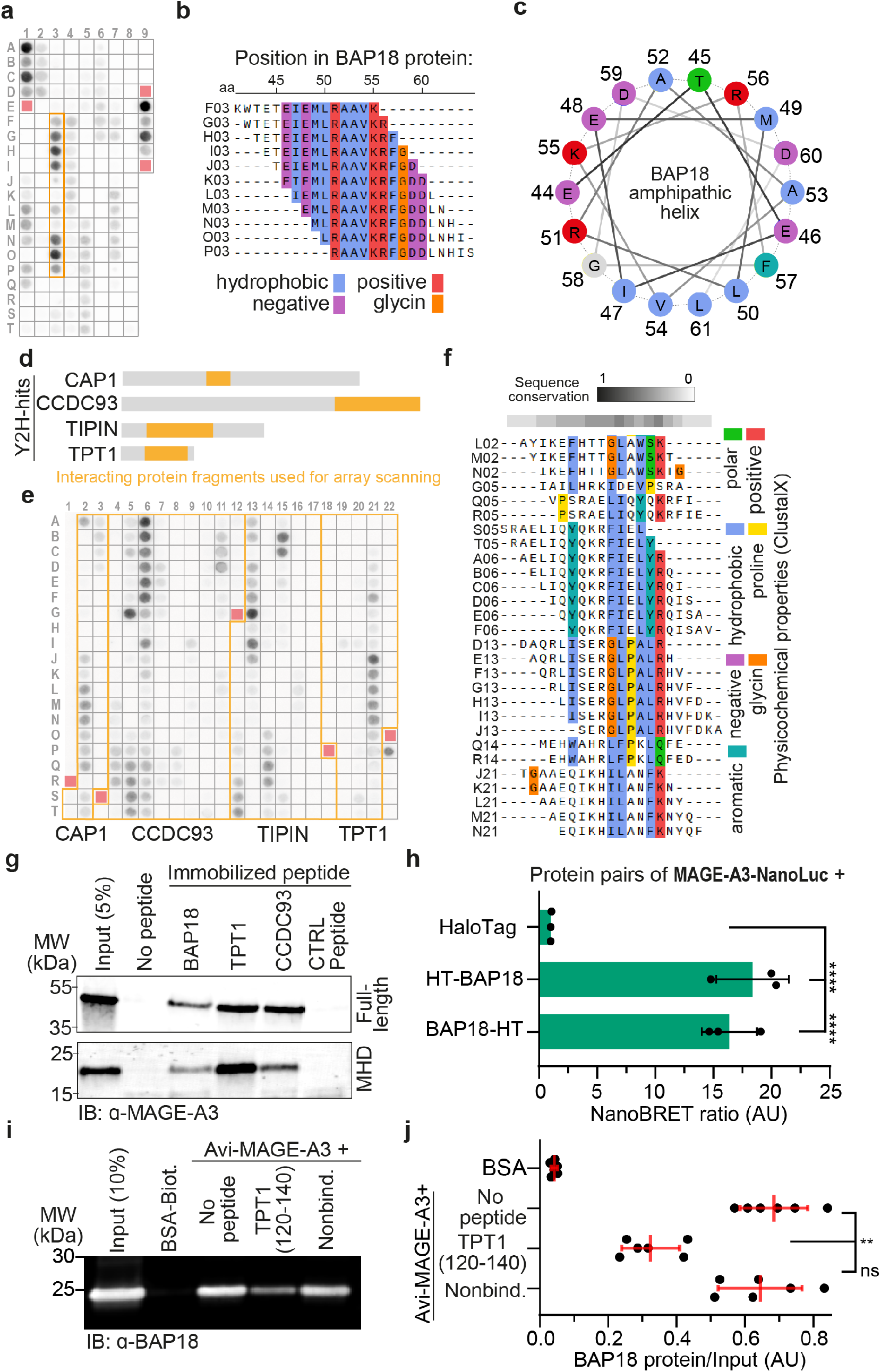
Identification of minimal MAGE-A3 binding peptide and characterization of BAP18/ MAGE-A3 interaction. **a**, Peptide array analysis of BAP18. Overlapping peptides producing the highest continuous interacting fragment are outlined with an orange box. Red squares indicate empty reference positions and positions A01-D01 and E09-H09 contain technical controls. **b**, MUSCLE peptide sequence alignment of peptide cluster producing the strongest interaction with MAGE-A3 protein. Peptide array position as shown in a, and amino acid position in full-length BAP18 are indicated. Physicochemical properties are according to ClustalX scale. **c**, Helical wheel projection illustrating the amphipathic character of the identified MAGE-A3 interacting helix of BAP18. Amino acid positions in the full-length BAP18 protein are indicated. **d**, Protein fragment hits of the Y2H screen using MAGE-A3 MHD (amino acids 100-314) as bait. Identified interacting fragments are shown in orange, with the respective full protein sequence outlined as gray bar. **e**, Peptide array analysis of Y2H interacting protein fragments, similar to panel a. **f**, MUSCLE peptide sequence alignment of various peptides producing signal after incubation with full-length MAGE-A3 on the peptide array. Physicochemical properties according to ClustalX scale. **g**, Immunoblot analysis of in vitro binding assay of full-length MAGE-A3 or MAGE-A3 MHD to biotinylated peptides immobilized on streptavidin beads. Peptides used for the in vitro binding assay encompass 20mer core interacting peptides as shown in b, f, derived from the indicated proteins. **h**, Quantification of NanoBRET cellular engagement assay between MAGE-A3-NanoLuc and N- or C-terminally HaloTAG-BAP18 protein, compared to HaloTAG. Significance was tested using Dunnett’s multiple comparisons test (*P* = 4.59^-6^ (HT-BAP18) *P* = 4.10^-5^ (BAP18-HT)). **i**, Fluorescent immunoblot analysis of in vitro binding assay of Avi-tagged, biotinylated and immobilized MAGE-A3 with full-length BAP18 protein. Avi-MAGE-A3 and BAP18 were incubated in the presence of the indicated peptides, compared to immobilized BSA. **j**, Quantification of in vitro binding assay as shown in i. BAP18 protein levels were normalized to the input. Significance was tested using a two-tailed Mann-Whitney U test (*P* = 2.16 ^-3^(+TPT1 peptide), *P* = 0.48 (+non-binding peptide). Biological replicates: *n* = 2 (**g**), *n* = 3 (**h**), *n* = 6 (**i, j**). Technical replicates: *n* = 1-2.

**Extended Data Figure 6.**
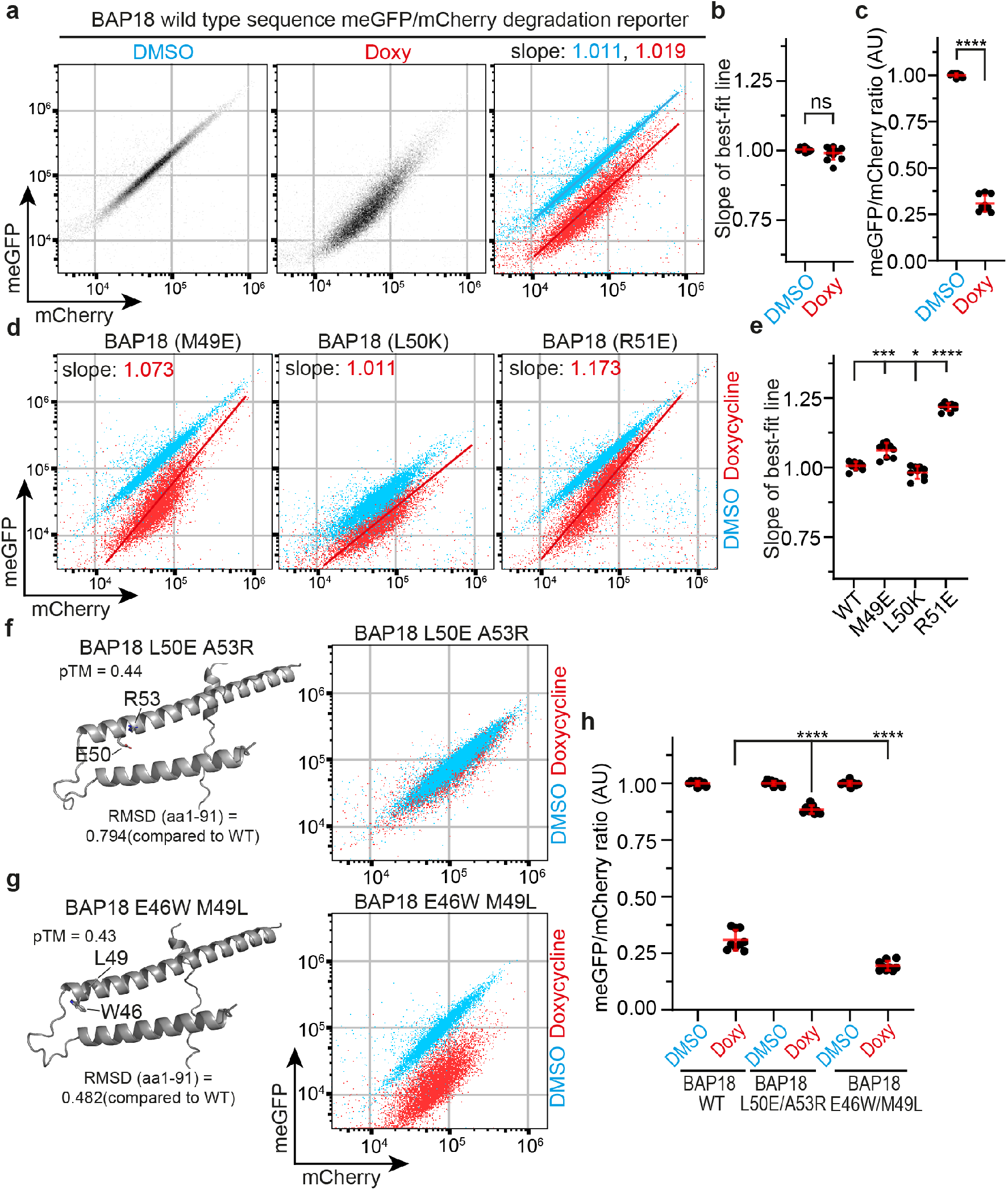
Investigation of BAP18 mutations modulating MAGE-A3 binding and degradation. **a**, FACS plots showing the levels of BAP18-meGFP/mCherry reporter stably expressed in iMAGE-A3-DLD-1 cells after DMSO or doxycycline treatment. **b**, Quantification of slopes of linear regression models for degradation reporter as shown in a. Bars indicate mean ± s.d. Significance was tested using a two-tailed Mann-Whitney *U*-test (*P* = 0.48). **c**, Quantification of meGFP/mCherry ratios for DMSO or doxycycline treatment in iMAGE-A3-DLD-1 cells as shown in a. and related to Fig 3b, d. Bars indicate mean ± s.d. Significance was tested using a two-tailed Mann-Whitney *U*-test (*P* = 4.11^-5^). **d**, FACS plots showing BAP18 single amino acid mutant reporter constructs stably expressed in iMAGE-A3-DLD-1 cells after DMSO or doxycycline treatment. **e**, Quantification of slopes of linear regression models for degradation reporters as shown in d. Bars indicate mean ± s.d. Significance was tested using a two-tailed Mann-Whitney *U*-test (*P* = 1.65^-4^ (M49E); *P* = 1.74^-2^ (L50K); *P* = 4.11^-5^ (R51E)). **f, g**, (left) BAP18 protein structure model, indicating amino acid positions of the indicated mutations. (right) FACS plot of BAP18 reporter carrying the indicated mutations stably expressed in iMAGE-A3 DLD-1 cells after DMSO or doxycycline treatment. **h**, Quantification of mutant reporter constructs stably expressed in iMAGE-A3 DLD-1 cells after DMSO or doxycycline treatment as shown in f, g. Bars indicate mean ± s.d. Significance was tested using a two-tailed Mann-Whitney *U*-test (*P* = 4.11^-5^ (L50E A53E, doxy); *P* = 4.11^-5^ (E46W M49L, doxy)). Biological replicates: *n* = 9 (**a**-**h**).

**Extended Data Figure 7.**
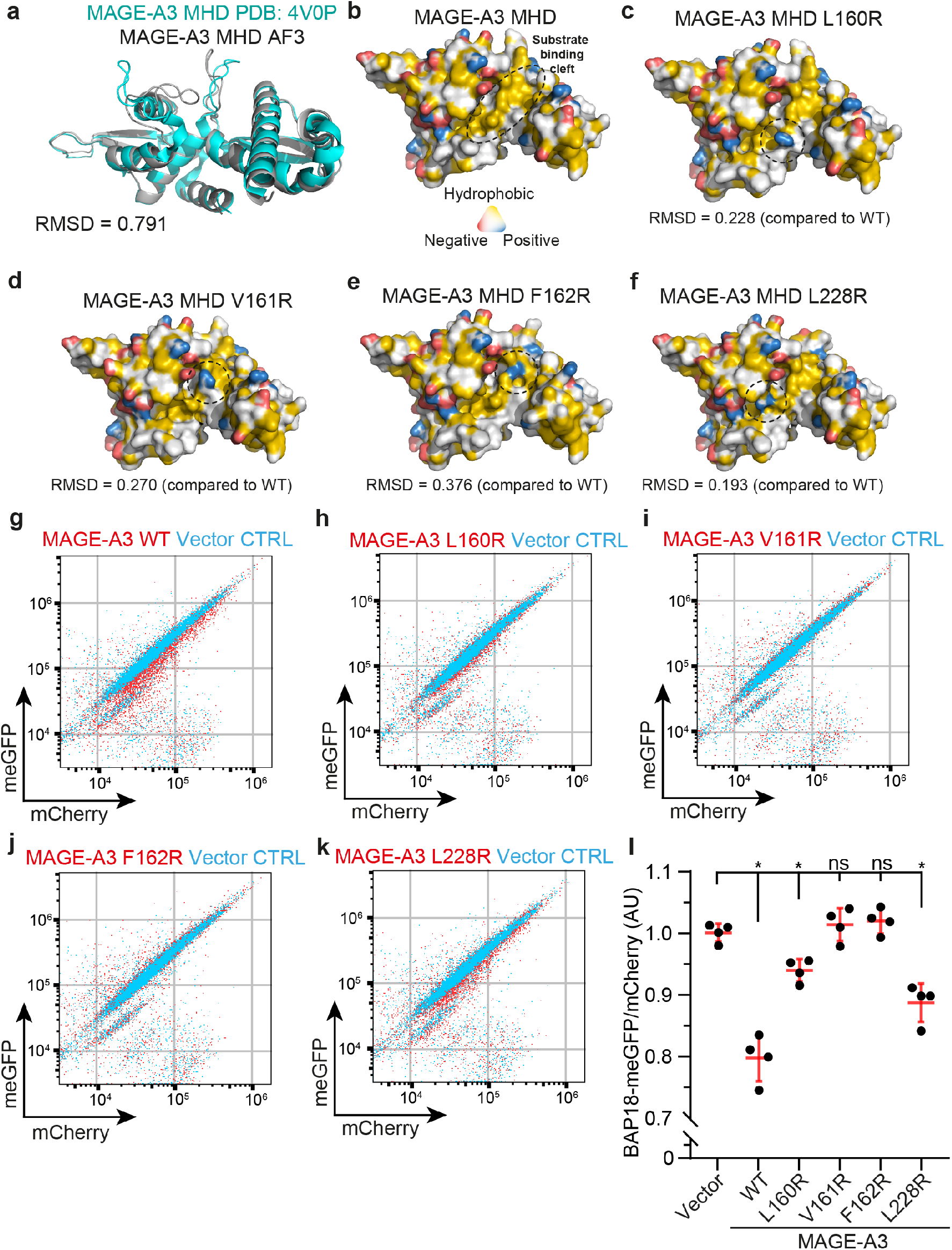
Investigation of MAGE-A3 mutations modulating BAP18 binding and degradation. **a**, Alignment of MAGE-A3 MHD from PDB 4V0P and AF3 prediction of MAGE-A3 MHD (amino acids 100-314). **b-f**, Surface representation of AF3 prediction of wild-type or mutant MAGE-A3 MHD, with YRB false color indicating charge and hydrophobicity. Hydrophobic, shallow substrate binding cleft (SBC) and the sites of change are indicated by a dashed outline. **g-k**, FACS plots of BAP18-meGFP/mCherry stably expressed in DLD-1 cells after transient overexpression of wild-type or mutant MAGE-A3 or vector control. **l**, Quantification of BAP18 reporter stably expressed in DLD-1 cells after transient overexpression of indicated MAGE-A3 constructs or vector for 48 hours. Bars indicate mean ± s.d. Significance was tested using a two-tailed Mann-Whitney *U*-test (*P* = 2.86^-2^ (WT); *P* = 2.86^-2^ (L160R); *P* = 0.69 (V161R); *P* = 0.20 (F162R); *P* = 2.86^-2^ (L228R)). Biological replicates: *n* = 4 (**g-l**). Technical replicates: *n* = 1-2.

**Extended Data Figure 8.**
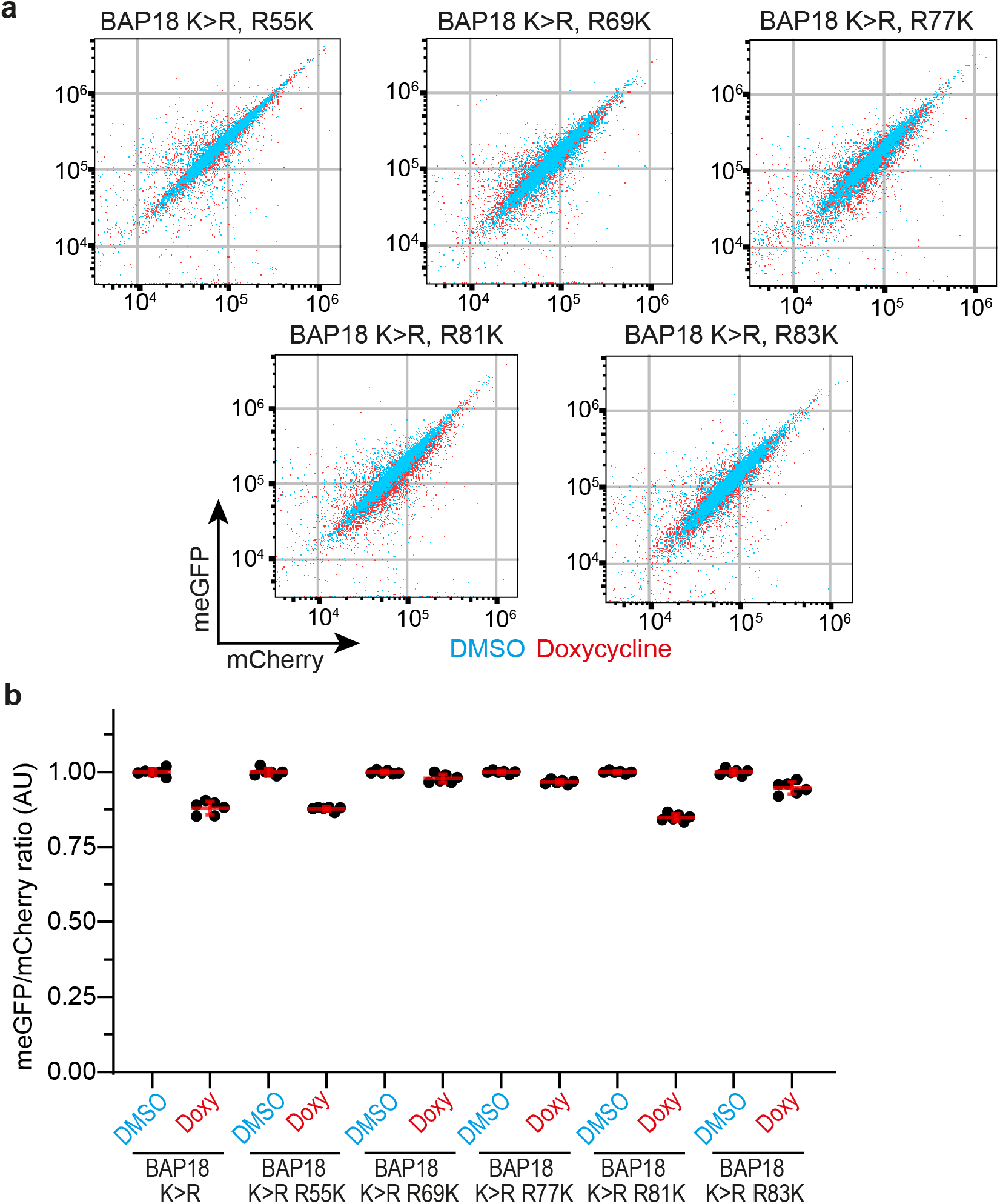
Effect of additional arginine-to-lysine reversion in BAP18 K>R reporter on MAGE-A3 induced degradation. **a**, FACS plots for BAP18 reporter constructs carrying the indicated mutations stably expressed in DLD-1-iMAGE-A3 cells after DMSO or doxycycline treatment for 48 hours. **b**, Quantification of panel a. Bars indicate mean ± s.d. Biological replicates: *n* = 6 (**a, b**).

**Extended Data Figure 9.**
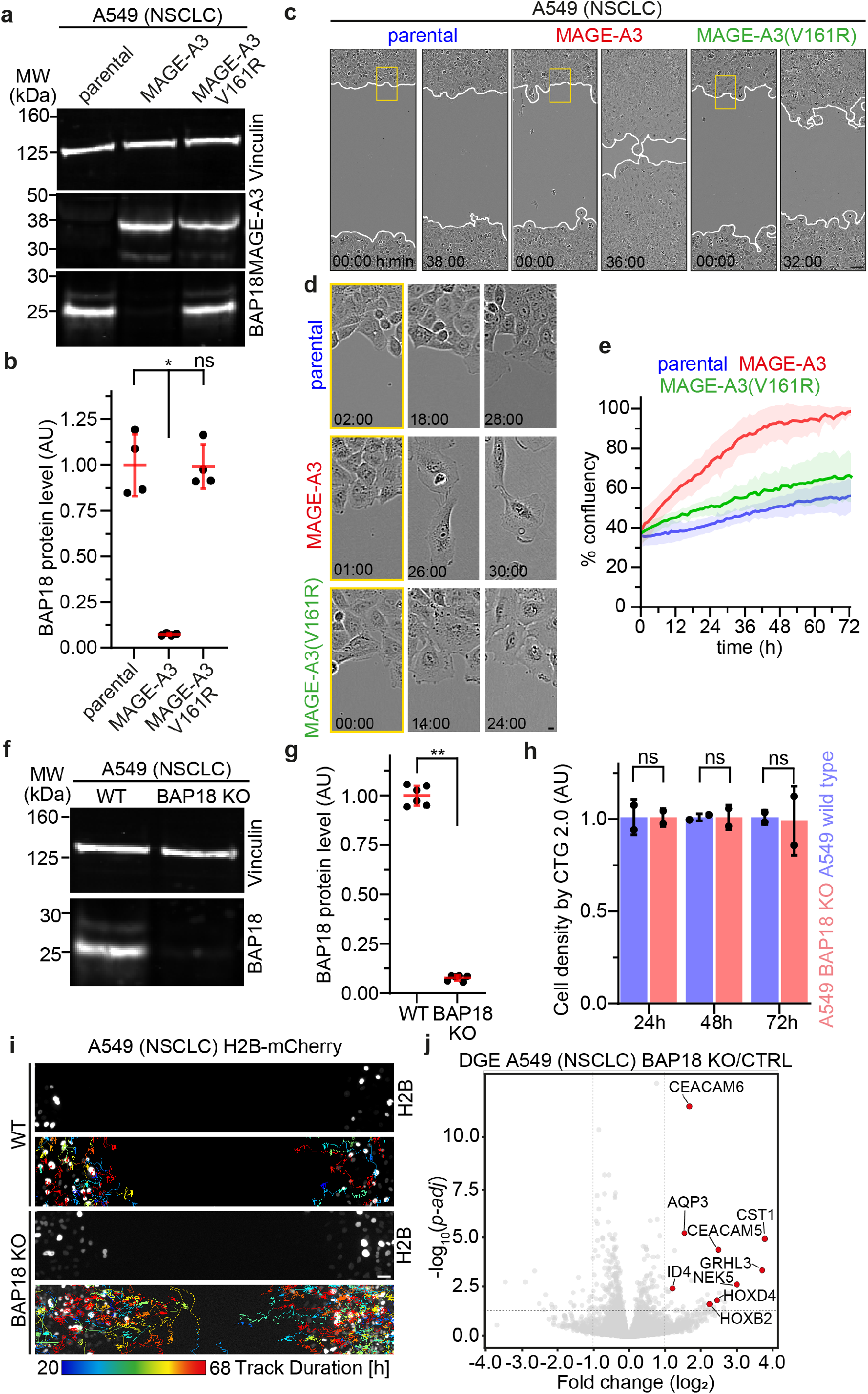
Characterization of MAGE-A3 overexpression and BAP18 KO in A549 (LUAD, NSCLC) cells. **a**, Fluorescent immunoblot analysis of MAGE-A3 or MAGE-A3(V161R) and BAP18 levels in stably transduced A549 cells compared to parental. **b**, Quantification of BAP18 protein levels in fluorescent immunoblot analysis as shown in a. Bars indicate mean ± s.d. Significance was tested using a two-tailed Mann-Whitney *U*-test (*P* = 2.86^-2^ (MAGE-A3); *P* = 0.89 (MAGE-A3 V161R)). **c**, Representative images of a wound healing assay of A549 cells stably expressing MAGE-A3 or MAGE-A3(V161R) compared to parental. Yellow rectangles indicate ROIs of insets shown in panel d. **d**, Pictures showing cell morphology at the boundary of the cell-free gap as shown in panel c. Time is shown as h:min. **e**, Quantification of cell migration into the cell-free gap shown in panel c. Data are mean ± s.d. **f**, Fluorescent immunoblot analysis of BAP18 protein levels in A549 BAP18 KO cells compared to wild type. **g**, Quantification of BAP18 protein shown in f. Bars indicate mean ± s.d. Significance was tested using a two-tailed Mann-Whitney *U*-test (*P* = 2.16^-3^). **h**, Quantification of cell proliferation by CTG measurement at 24, 48 or 72 hours post cell seeding for A549 BAP18 KO cells compared to wild-type. Values are normalized to mean of wild type for each measurement timepoint. Bars indicate mean ± s.d. Significance was tested using a two-tailed Mann-Whitney *U*-test (*P* = 0.99 (24h, 48h, 72h)). **i**, Single cell tracks of H2B-mCherry labelled A549 cells. Track color indicates track duration, depicted tracks filtered by >19h duration. **j**, Volcano plot displaying DEGs between A549 BAP18 KO and wild type cells. Significance and fc-thresholds are indicated as dashed gray lines (FDR-adjusted *P*-values < 0.05; |log2FC| > 1). Selected highly upregulated genes are labelled red. Biological replicates: *n* = 4 (**a, b**), *n* = 6 (**d, e**), *n* = 2 (**f**), *n* = 3 (**g**). Technical replicates: *n* = 2-4. Scale bars, 1 µm (**c**), 100 µm (**g**).

**Extended Data Figure 10.**
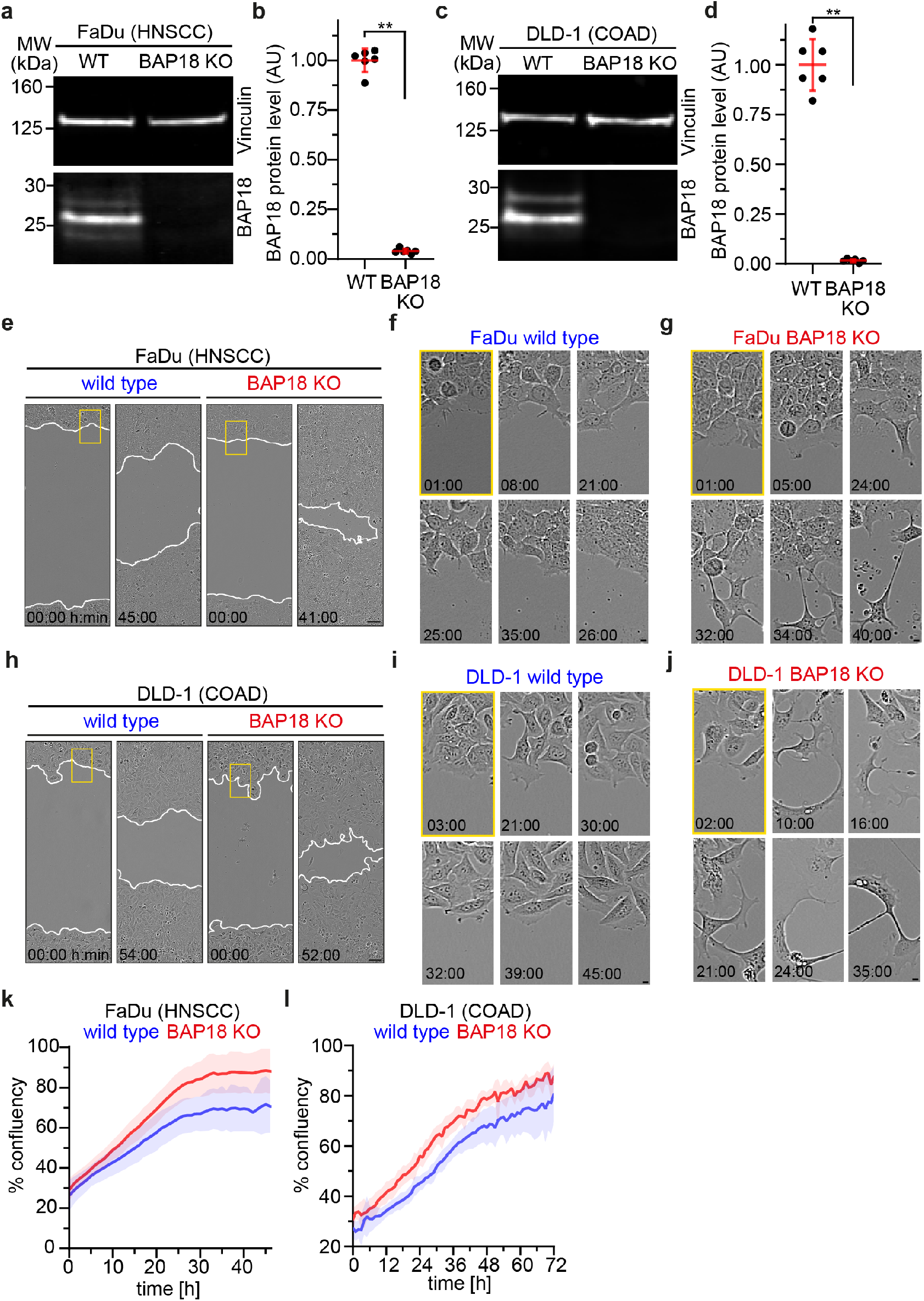
Characterization of BAP18 KO in FaDu (HNSCC) and DLD-1 (COAD) cells. **a**, Fluorescent immunoblot analysis of BAP18 protein levels in FaDu BAP18 KO cells compared to wild-type. **b**, Quantification of BAP18 protein levels in fluorescent immunoblot analysis as shown in a. Bars indicate mean ± s.d. Significance was tested using a two-tailed Mann-Whitney *U*-test (*P* = 2.16^-3^). **c**, Fluorescent immunoblot analysis of BAP18 protein levels in DLD-1 BAP18 KO cells compared to wild-type. **d**, Quantification of BAP18 protein levels in fluorescent immunoblot analysis as shown in c. Bars indicate mean ± s.d. Significance was tested using a two-tailed Mann-Whitney *U*-test (*P* = 2.16^-3^). **e**, Migration assay of FaDu cells after stable CRISPR/Cas9 induced KO of BAP18 compared to wild-type. Yellow rectangles indicate ROIs of insets shown in panels f, g. **f**, Insets of wild type FaDu cell morphology at the border of the cell-free gap as shown in e. **g**, Insets of BAP18 KO FaDu cell morphology at the border of the cell-free gap as shown in e. **h**, Migration assay of DLD-1 cells after stable CRISPR/Cas9 induced KO of BAP18 compared to wild-type. Yellow rectangles indicate ROIs of insets shown in panels i, j. **i**, Insets of wild type DLD-1 cell morphology at the border of the cell-free gap as shown in h. **j**, Insets of BAP18 KO DLD-1 cell morphology at the border of the cell-free gap as shown in h. **k**, Quantification of cell migration into the cell-free gap of FaDu BAP18 KO cells compared to wild-type as shown in e. Data are mean ± s.d. **l**, Quantification of cell migration into the cell-free gap of DLD-1 BAP18 KO cells compared to wild-type as shown in h. Data are mean ± s.d. Biological replicates: *n* = 6 (**a-d**), *n* = *3* (**e-l**). Scale bars, 100 µm (**d**), 1 µm (**e, f**). Time is shown as h:min. (**d**-**f**).

