## Supplementary Data Fig 1 for "Oncogenic E3-ligase adaptors MAGE-A3/6 promote cancer cell migration via BAP18 degradation"

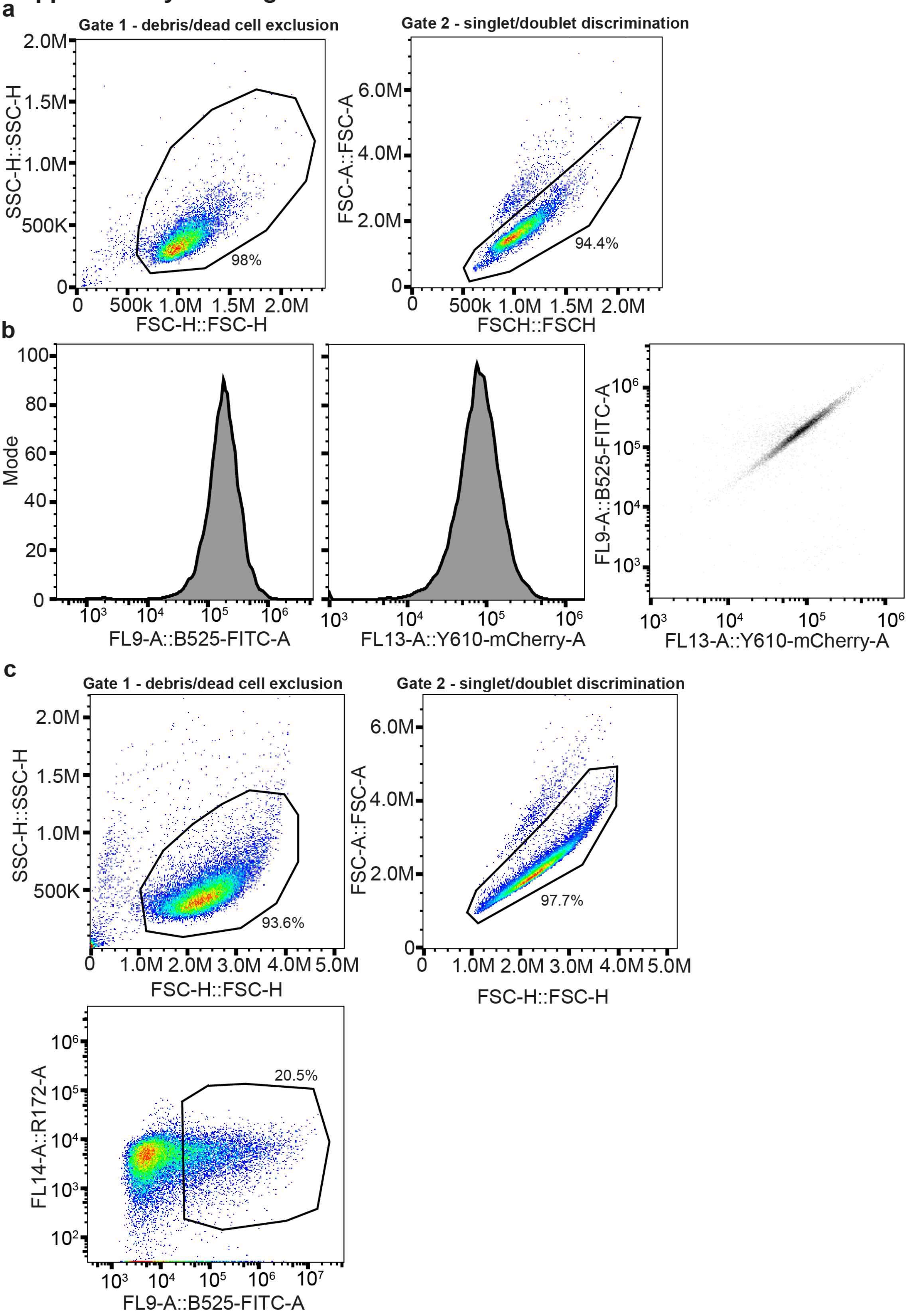


**Supplementary Data Figure 1. Gating strategy for analysis of BAP18-meGFP/mCherry reporter cells and sorting of Cas9-TagBFP2 transfected cells**

**a**, Representative gating strategy for all reporter assay utilizing BAP18-meGFP/mCherry reporter cell lines. The cells stably expressing the reporter constructs were in addition transfected or treated with doxycycline (iMAGE-A3-DLD-1 cells). **b**, Live cell singlets were used to measure meGFP and mCherry histograms, of which the medians were used for quantifying relative BAP18 downregulation. A representative meGFP over mCherry scatterplot is shown for a control DMSO treated sample. **c**, Representative gating strategy for sorting all-in-one Cas9-TagBFP2-dgRNA transfected DLD-1-iMAGE-A3 cells to induce TRIM28KO. Live cell singlets belonging to the top 20.5% BFP positive population were single-cell sorted into 96-well plates to recover clonal cells. Each plot shows the gated subpopulation for the preceding plot, indicating the respective fractions.
